# Early-life exposure to commensal lung bacteria primes innate antiviral immunity and prevents RSV immunopathology in neonatal mice

**DOI:** 10.64898/2025.12.05.692516

**Authors:** Quentin Marquant, Claire Chottin, Cécile Ferret, Eloi Poucet, Vinciane Saint-Criq, Daphné Laubreton, Carole Drajac, Edwige Bouguyon, Aude Remot, Samuel Constant, Song Huang, Edouard Sage, Isabelle Schwartz-Cornil, Muriel Thomas, Sabine Riffault, Delphyne Descamps

## Abstract

The lung mucosa during the neonatal period and early infancy is especially susceptible to respiratory syncytial virus (RSV) infection. Respiratory disease severity is strongly influenced by the age at first exposure, which may also affect the trajectory of airway function. Early life represents a critical window for lung development and initial microbiota colonization, both of which shape mucosal immune responses to RSV. The contribution of early lung-colonizing bacterial strains to the establishment of the lung microbiota and susceptibility to RSV infection in neonatal mice remains poorly characterized. In the present study, we showed that early-life primo-colonizing bacterial strains in the mouse lung differentially induce innate immune responses and influence RSV susceptibility in *ex vivo* models using lung explants and alveolar macrophages (AMs). We identified a specific bacterial strain (strain 17) whose prior exposure enhances type I interferon (IFN-I) responses in AMs upon RSV infection and reduces viral replication both *ex vivo* and *in vivo* in lung tissues. Intranasal administration of this strain during early life prevented the development of immunopathological responses upon RSV reinfection in adult mice. Finally, using a translational human airway epithelium model, we demonstrated that pre-exposure to strain 17 restricts RSV spread without cytotoxicity, likely *via* enhanced β-defensin 2 production. These findings highlight the potential of early-life microbiota modulation as a promising intervention for preventing RSV disease and its long-term respiratory consequences.

**Author summary:** Respiratory syncytial virus (RSV) is a major cause of severe respiratory infections in newborns and infants, a period during which the lungs and immune system are still maturing. Simultaneously, the respiratory tract undergoes its first microbial colonization by so-called primo-colonizing bacteria — the first bacterial species to establish themselves on neonatal lower respiratory mucosal surfaces. These early microbial pioneers may play an unsuspected role in shaping how the immature immune system responds to viral threats. In this study, we investigated whether these early lung-colonizing strains influence RSV susceptibility. Using neonatal mouse models, lung explants, and alveolar macrophages, we found that distinct bacterial strains trigger markedly different innate immune responses. Prior exposure to one strain in particular, strain 17, was associated with enhanced type I interferon responses, reduced viral replication, and protection against harmful inflammation upon RSV reinfection later in life. In a human airway epithelium model, pre-exposure to strain 17 also restricted RSV spread without tissue damage, potentially through increased β-defensin 2 production. These findings provide proof-of-concept that the bacterial strains establishing in the neonatal lung can influence future immune reactivity and antiviral responses, and suggest that early-life respiratory microbiota modulation could represent a promising strategy to prevent severe RSV disease.

## Introduction

The respiratory syncytial virus (RSV), known as the primary cause of bronchiolitis in infants, is a major contributor to lower respiratory tract infections (LRTIs), and remains the leading cause of child mortality worldwide [1]. Severe early-life RSV infection has notably been associated with recurrent LRTIs, wheezing, asthma, impaired lung function, and chronic respiratory disease later in life [2]. New passive immunisation strategies, such as maternal vaccination or administration of monoclonal antibodies, have been approved to reduce the burden of acute RSV infection in high-risk infants. However, some complications have been reported, including an increase in preterm births [3]. Direct vaccination in children remains a challenge, particularly after the FDA recently suspended all RSV vaccine trials in infants following reports of increased cases of severe illness in one trial [4]. The mechanisms underlying pulmonary hypersensitivity to RSV in early-life and its long-term health effects remain to be elucidated in order to develop effective prophylactic strategies. An increasing number of studies on RSV-infected neonatal mice (less than 7 days old at initial infection) have validated this model as relevant for understanding RSV infection in human infants [5]. It has been shown that the age at first infection plays a key role in shaping subsequent immune responses, notably by inducing epigenetic modifications [6]. This may increase the severity of later RSV infections [7], potentially by driving a type 2-biased immunity [8], which promotes allergic responses later in life [6]. Moreover, the early-life in mice is characterized by an immediate permissiveness to RSV replication, which gradually decreases with age as the ability to mount type I interferon (IFN-I) responses improves [9]. IFN-I are rarely detected in nasal washes from RSV-infected children [10]. However, IFN-α and IFN-β control viral pathogenesis by inducing interferon-stimulated genes that provide an early antiviral defense and shape inflammatory and adaptive immune responses [11]. IFN-I responses appear to play a key role in the outcome of RSV infection [11], as treatments that enhance IFN-I levels in infected neonatal mice have been shown to reduce viral replication and long-term RSV-induced lung effects [12, 13]. Thus, modulation of pulmonary innate immune defenses during the perinatal period can affect susceptibility to RSV infection, and determine long-term respiratory outcomes.

Early life, spanning the neonatal period and early infancy, is a critical window during which the pulmonary immune system matures alongside the establishment of the respiratory microbiome, while the lungs themselves continue to develop. Disturbances during this period may have lasting effects on respiratory health [14]. When the balance between the three interconnected pulmonary components (namely the airway epithelium, immune cells, and resident microbiota) is disrupted, it can ultimately increase susceptibility to infections and predispose individuals to chronic respiratory diseases. [15]. There is increasing evidence that postnatal lung mucosa is biased toward a type 2 immune response, as indicated by the spontaneous accumulation of type 2 innate lymphoid cells (ILC2), eosinophils, and basophils, along with the production of cytokines such as IL-33 and IL-13 [16], which are typically involved in anti-inflammatory responses and tissue repair. Such a tissue environment is prone to the early polarization of tissue-resident alveolar macrophages (AMs) toward a tolerogenic, anti-inflammatory M2 phenotype [17]. Moreover, the lung microbiota establishes rapidly after birth and it shaped by factors linked to the host and the environment [18–20]. A recent prospective pediatric cohort study showed that rapid changes in the upper airway bacterial microbiota between birth and 3 months of age are accompanied by dynamic maturation of JAK/STAT and antiviral immune pathways, including a marked increase in interferon-related signaling during the first year of life, highlighting a close link between bacterial communities and early-life respiratory immunity [21]. In addition, the lung microbiota contributes to immune tolerance in early life according to an asthma model [18] and attenuates innate immune reactivity to PAMPs [22].

The incidence of respiratory infections and the occurrence of dysbiosis in the lung microbiota are closely related [23]. The severity of RSV infection, [24, 25] and the persistence of symptoms in children may be influenced by the bacterial ecosystem [26]. Conversely, the presence of *Lactobacillus* in the nasopharynx is associated with a reduced risk of childhood wheezing illnesses at 2 years of age [27]. In mice, viable commensal bacteria are detected in the lungs as early as 3 days of age, with their number and diversity increasing over time [18] and reaching a plateau by weaning at around 21 days of age [20]. A collection of primo-colonizing bacteria, defined as the first bacterial species that establish on neonatal lower respiratory mucosal surfaces, was established through their isolation from lung homogenates of neonatal mice using aerobic or anaerobic culture conditions [20]. Intranasal administration of certain strains of this collection during early life influenced the outcome of an allergic asthma model [20], while instillation of other strains in adult mice attenuated pulmonary inflammation following *Mycobacterium tuberculosis* infection [28]. These findings suggest that lung primo-colonizing commensal bacteria can modulate pulmonary immunity and influence susceptibility to respiratory diseases. However, the mechanisms by which early bacterial colonization of the lungs modulates immune responses and determines both immediate and long-term respiratory outcomes, including during early-life RSV infection, remain largely unexplored [29]. In the present study, we provide evidence that early-life commensal lung bacteria can contribute to shaping immune responses and influencing RSV susceptibility in neonatal mice. We established a proof of concept demonstrating that it is possible to select a single strain from a panel of lung primo-colonizing commensal bacteria based on its immunomodulatory and antiviral properties on the neonatal lungs. One bacterial strain in particular, hereafter referred to as strain 17, promoted IFN-I antiviral pathways and reduced RSV replication both *ex vivo* and *in vivo* in lung tissues. Finally, its intranasal administration to neonatal mice prevented early-life immunopathological priming, which became apparent after a second exposure to RSV in adulthood.

## Results

### Primo-colonizing bacteria induce distinct innate immune activation patterns in neonatal lung mucosa *ex vivo* model

The goal was to test the concept that a commensal and primo-colonizing strain can promote a protective type 1 immune response in the lungs during early life, associated with improved control of RSV infection. To evaluate the immunostimulatory effects of primo-colonizing bacterial strains (S1 Table) on the neonatal lung, we established a PCLS-based screening assay using lung tissue from 6-day-old mice, an *ex vivo* model that recapitulates pulmonary cytokine responses [30]. PCLS were exposed to each strain at MOI=50 (Fig 1A). Strains were standardized at the onset of the stationary phase to ensure a comparable bacterial physiological state at the time of stimulation, while acknowledging that some variability in bacterial growth or biomass during the 16 h incubation cannot be entirely excluded. Among 20 strains tested, two (strains 1 and 11) caused significant cytotoxicity after 16h and were excluded, while most showed cytotoxicity levels comparable to the control medium (mean = 21.3% ± 1.1%), indicating no harmful effects (S1A Figure). Cytokines were quantified using a 15-plex assay and grouped by functional category (pro-inflammatory, type 1 immunity, type 2 immunity, Th17/Th22/Th9-responses, and lymphocyte proliferation, S1B-F Fig). Both the nature and magnitude of cytokine production by PCLS varied across bacterial strains. Strains 2, 19 and 16 induced high levels of pro-inflammatory cytokines such as TNF-α and IL-6, together with robust a type 2 immune response, including IL-10 (S1B and S1D Fig). In contrast, most other strains induced low cytokine production. Thus, strain 17, identified as *Escherichia coli* by 16S sequencing and mass spectrometry (S1 Table), also induced type 2 cytokines (S1D Fig) but was unique in triggering a strong type 1 cytokine profile, characterized by high production of IFN-γ and IL-12p70 (Fig 1B), along with increased IL-22 and IL-9 secretion (S1E Fig). To highlight strain-specific immune signatures, cytokine levels were expressed as fold change relative to control PCLS and visualized as heat map (Fig 1C), with hierarchical clustering revealing groups of strains with similar immunostimulatory patterns. Overall, primo-colonizing lung bacteria displayed distinct immunostimulatory capacities in PCLS from neonatal mice, with strain 17 standing out for its pronounced type 1 cytokine response and lack of cytotoxicity.

**Figure. 1:**
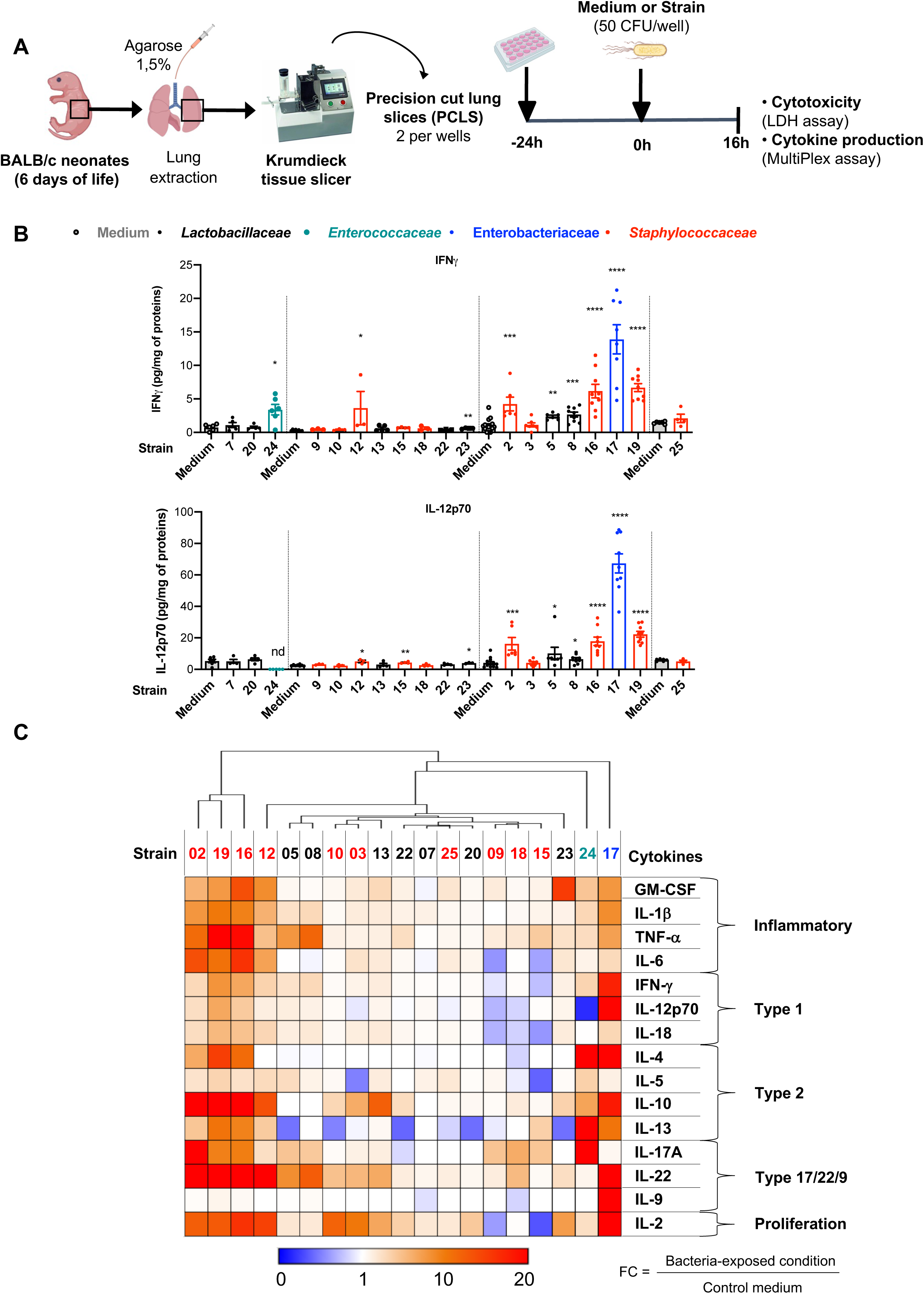
Cytokine production by precision-cut lung slices (PCLS) from neonatal mice varies in response to exposure to lung primo-colonizing bacteria. **A**. Experimental design. The lungs from 6-day-old mice were isolated and cut with a Krumdieck tissue slicer to obtain PCLS. They were next exposed to bacteria (50 CFU/well) or cultured in medium control. Supernatants were collected 16h after stimulation with bacterial strains and PCLS were lysed. **B**. Secretion of cytokines associated with type 1 immune response (IFN-β, IL-12p70). Results are expressed as mean ± SEM and represent a pool of five independent experiments (n = 2-5 biological replicate samples/condition). \**P* < 0.05, \*\**P* < 0.01, \*\*\**P* < 0.001 and \*\*\*\**P* < 0.0001 between the control medium and the strain-exposed condition (ANOVA, Kruskal-Wallis test followed by Dunn’s post-hoc test). **C**. The heat map illustrates the fold-change (FC) of cytokine production induced after bacterial exposure. For each strain and cytokine secretion, the FC were calculated using the following formula: FC = (average of the stimulated conditions)/(average of the control conditions). FC were entered into Morpheus online software (Broad Institute) to obtain the heatmaps.

### Innate immune activation by strain 17 is associated with reduced RSV replication *ex vivo*

We next focused on the early events following RSV infection to explore how strain 17 affects early innate antiviral response. To assess whether strain 17 influences early-life RSV susceptibility, we performed co-exposure experiments using PCLS from 6-day-old mice. PCLS were first exposed to strain 17 (50 CFU/well as previously used [20]) to elicit its immunostimulatory effects, and then infected with luciferase-expressing RSV (RSV-Luc) or mock-treated for 16h to evaluate viral replication (Fig 2A). Strains 16, 24, and 25, displaying distinct cytokine profiles (see S1 Fig), were included as comparators. Cytotoxicity induced by these strains, RSV-Luc, or their combination did not exceed 21% and was comparable to that of the control medium. Thus, co-exposure did not affect PCLS viability and supporting the safe use of these bacteria during RSV infection (Fig 2B). Viral replication was quantified in PCLS lysates using a luciferase activity assay (Fig 2C). PCLS exposed to strain 25 showed a slight increase in luciferase activity compared to the control condition (not statistically different), suggesting a potential enhancement of RSV replication associated with the presence of strain 25. Pre-exposure to strains 16 or 24 resulted in only a slight reduction in RSV replication (non stastistically significant). In contrast, exposure to strain 17 resulted in a strong significant reduction in luciferase activity compared to both control and PCLS exposed to strain 16 or 24 conditions. Altogether, these results suggest that pre-exposure to lung primo-colonizing commensal bacteria can modulate RSV replication in PCLS from 6-day-old mice, with strain 17 emerging as the most effective strain in reducing RSV replication.

**Figure. 2:**
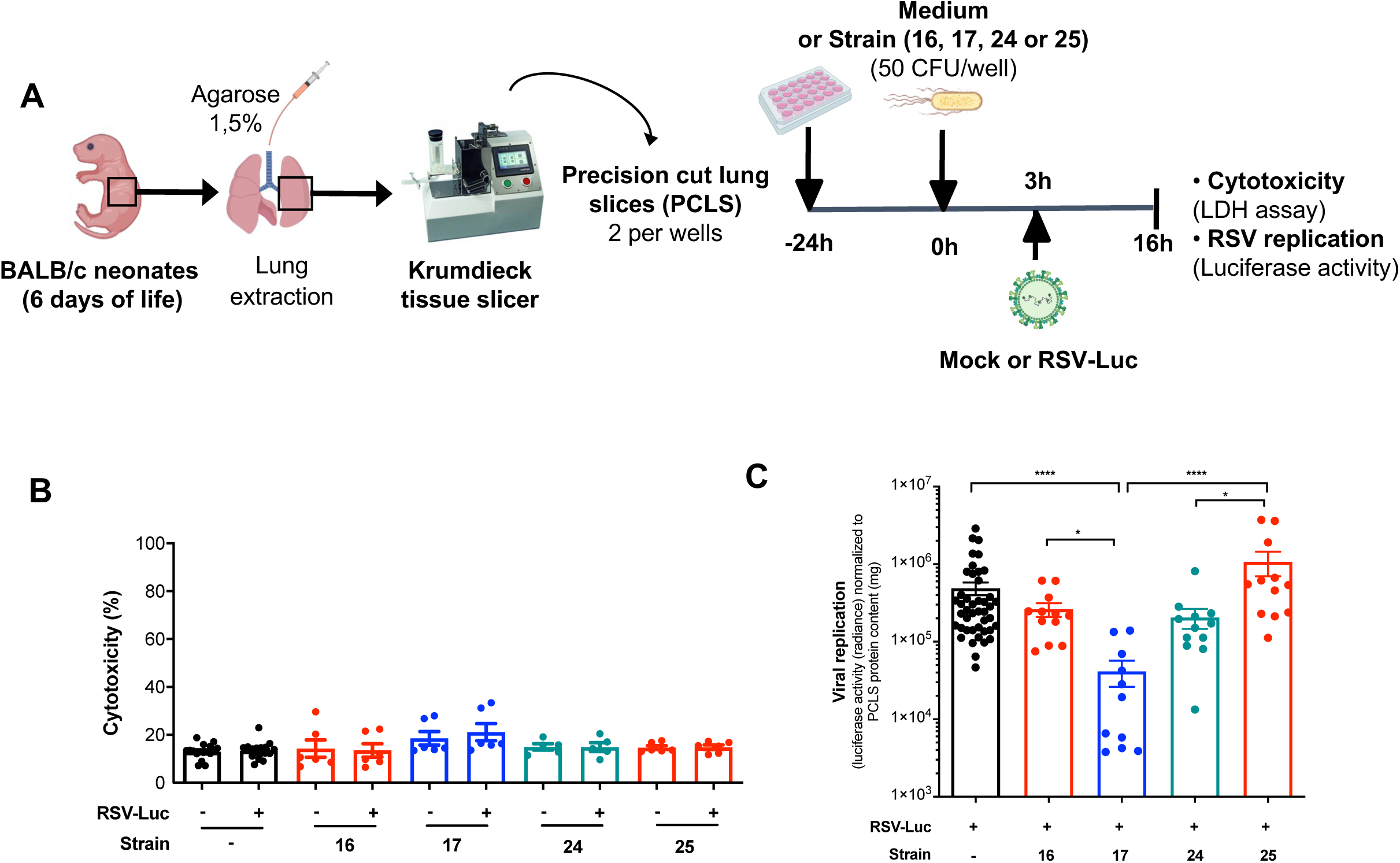
Exposure to primo-colonizing commensal bacteria differentially affects RSV replication in neonatal PCLS. **A**. Experimental design. The lungs from 6-day-old mice were isolated and cut with a Krumdieck tissue slicer to obtain PCLS. They were next exposed to strains 16, 17, 24 or 25 (50 CFU/well) or cultured in control medium. PCLS were infected with RSV expressing luciferase (RSV-Luc, 7.187 x 10^4^ PFU/well) or mock control (-) 3h after bacterial stimulation. Supernatants were collected 16h after bacterial stimulation and PCLS were lysed. **B**. Cytotoxicity observed in PCLS was measured by LDH activity assay. Results are expressed as the mean ± SEM and represent a pool of four independent experiments (n = 5-18 per group in simplicate). **C**. RSV-Luc replication was assessed by measuring luciferase activity in lysates of PCLS exposed or unexposed to strains. Results are expressed as mean ± SEM and represent a pool of four independent experiments (n = 6-21 per group in duplicate). \**P* < 0.05 and \*\*\*\**P* < 0.0001 between the control medium or the strain-exposed condition (ANOVA, Kruskal-Wallis test followed by Dunn’s post-hoc test).

We also investigated the effect of strain 17 on AMs, the main pulmonary resident immune cells, involved in IFN-I production during RSV infection [31]. AMs isolated from BAL of 6-day-old mice were first exposed to strain 17 (50 CFU/well as previously used [20]) or control medium, followed by exposure to RSV (Fig 3A). After 16h of incubation, exposure of AMs to strain 17 alone induce a slight but unsignificant production of IL-6, TNF-α, and IFN-I in the supernatants compared to basal control conditions (no strain, no RSV, Fig 3B-C). When AMs exposed to strain 17 were subsequently infected with RSV, they produced markedly more inflammatory (IL-6 and TNF-α, Fig 3B) and antiviral (IFN-α and IFN-β, Fig 3C) cytokines than RSV-infected AMs without bacterial pre-exposure. Notably, RSV-infected AMs produced 3-fold higher levels of IFN-I when exposed to strain 17. Thus, neonatal AMs exposure to strain 17 improved their innate immune activation, contributing to better control of RSV infection.

**Figure. 3:**
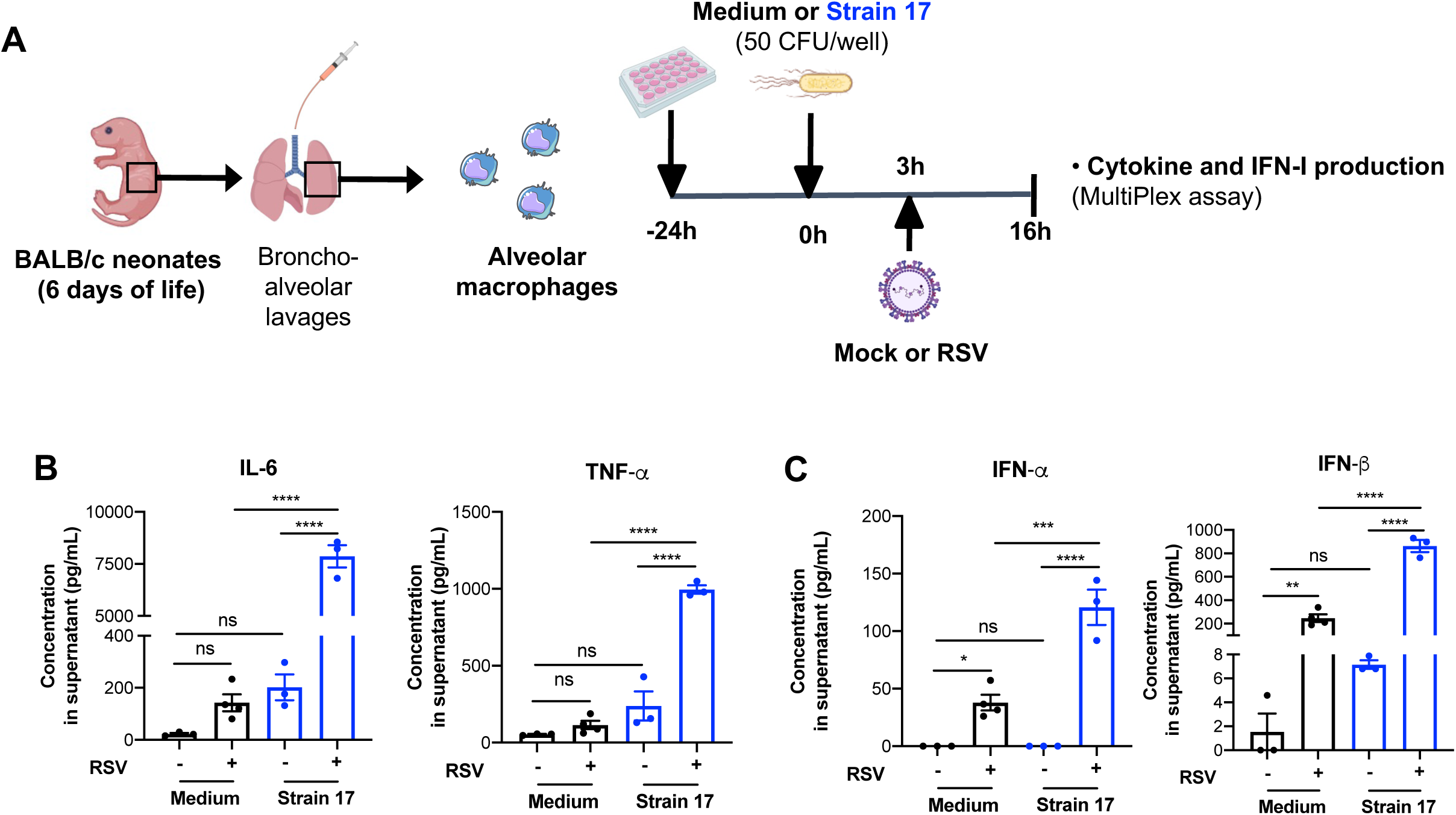
Cytokines and type I interferon (IFN-I) production by RSV-infected alveolar macrophages (AMs) from neonatal mice is enhanced in response to exposure to strain 17. **A**. Experimental design. AMs isolated from BAL of neonatal mice were exposed to strain 17 (50 CFU/well) or cultured in medium control. AMs were infected with rRSV-long (MOI = 5) 3h after bacterial stimulation. Supernatants were collected 16h after bacterial stimulation and AMs were lysed. The production of pro-inflammatory cytokines (IL-6 and TNF-α, **B**) and IFN-I (IFN-α and IFN-β, **C**) was measured in supernatants using multiplex immunoassay. Data are representative of two independent experiments. Means ± SEM are represented. \**P* < 0.05, \*\**P* < 0.01, \*\*\**P* < 0.001 and \*\*\*\**P* < 0.0001 (2way ANOVA with Tukey’s multiple comparisons test).

### Pulmonary exposure to strain 17 decreases neonatal susceptibility to RSV infection through innate immune activation

Collectively, these *in vitro* results identify strain 17 as non cytotoxic strain capable of enhancing IFN-I production, contributing to reduced RSV replication. We therefore evaluated whether pulmonary exposure to strain 17 confers protection against RSV infection in neonatal mice. To assess the *in vivo* relevance of its effects, we first investigated whether intranasal (i.n.) administration of this commensal strain (10^6^ CFU; dose and method previously shown to deliver sufficient bacteria to the lungs [20]) was non-pathogenic in neonatal mice. No mortality was observed in neonatal mice following administration of strain 17 (data not shown). A modest increase in neutrophil counts was detected in BALs, although total cell numbers remained unchanged S2 Fig). Next, we performed a dose-response study to determine the minimal amount of strain 17 required to achieve its *in vivo* immunomodulatory effect. Neonatal mice received three different amounts of strain 17 or PBS diluent intranasally at 3- and 5-days-old to modulate the neonatal lung mucosa before infection with RSV-Luc at day 6 (Fig 4A). Mice were euthanized 1 or 2 d.p.i. to assess viral replication in the upper and lower respiratory tracts, as well as the associated immune response in lung tissues. As previously shown [9, 13], i.n. RSV-Luc infection of neonatal mice led to rapid viral replication in the nasal cavity, reaching 1.42 × 10⁵ ± 0.38 × 10⁵ and 0.89 × 10⁵ ± 0.28 × 10⁵ photons/mg at 1 and 2 d.p.i., respectively (Fig 4B). In PBS-treated neonates, lung replication was detectable at 1 d.p.i. (1.02 × 10⁴ ± 0.38 × 10⁴ photons/mg) and increased by 2 d.p.i. (2.09 × 10⁴ ± 0.32 × 10⁴ photons/mg), confirmed by RSV *N gene* expression (Fig 4C-D). Strain 17 exposure was associated with significantly reduced nasal viral replication at 1 d.p.i. at all doses (3-4-fold, Fig 4B), whereas only the 10⁶ CFU dose was associated with significantly lower lung viral replication (0.098 × 10⁴ ± 0.047 × 10⁴ photons/mg, Fi 4C). The 10⁵ CFU dose showed delayed lung protection at 2 d.p.i., while the 10³ CFU dose had no effect, indicating that upper airway protection requires lower doses, whereas lung protection needs higher or delayed dosing (Fig 4B-D). RSV infection in neonatal mouse lungs induced low *IFN-I* gene expression at 1 d.p.i. (Fig 5A), and led to a slight, non-significant, increase in BAL cellularity between 1 and 2 d.p.i. (Fig 5B), primarily due to neutrophil recruitment (Fig 5D), as previously shown [9, 12, 13]. In neonates treated with high doses of strain 17, *IFN-I gene* expressions were increased at 1 d.p.i. (Fig 5A), while total BAL cells (Fig 5B) and macrophages (Fig 5C) remained unchanged. In contrast, neutrophil recruitment was significantly enhanced from 1 to 2 d.p.i., with an ∼8-fold increase at 2 d.p.i. in RSV-infected mice pre-treated with the highest doses of strain 17 (Fig 5D). Overall, these results demonstrate that neonatal exposure to strain 17 enhances early innate immune responses, leading to reduced RSV replication, dose-dependent protection of the lungs and a strong protective effect in the nasal cavity.

**Figure. 4:**
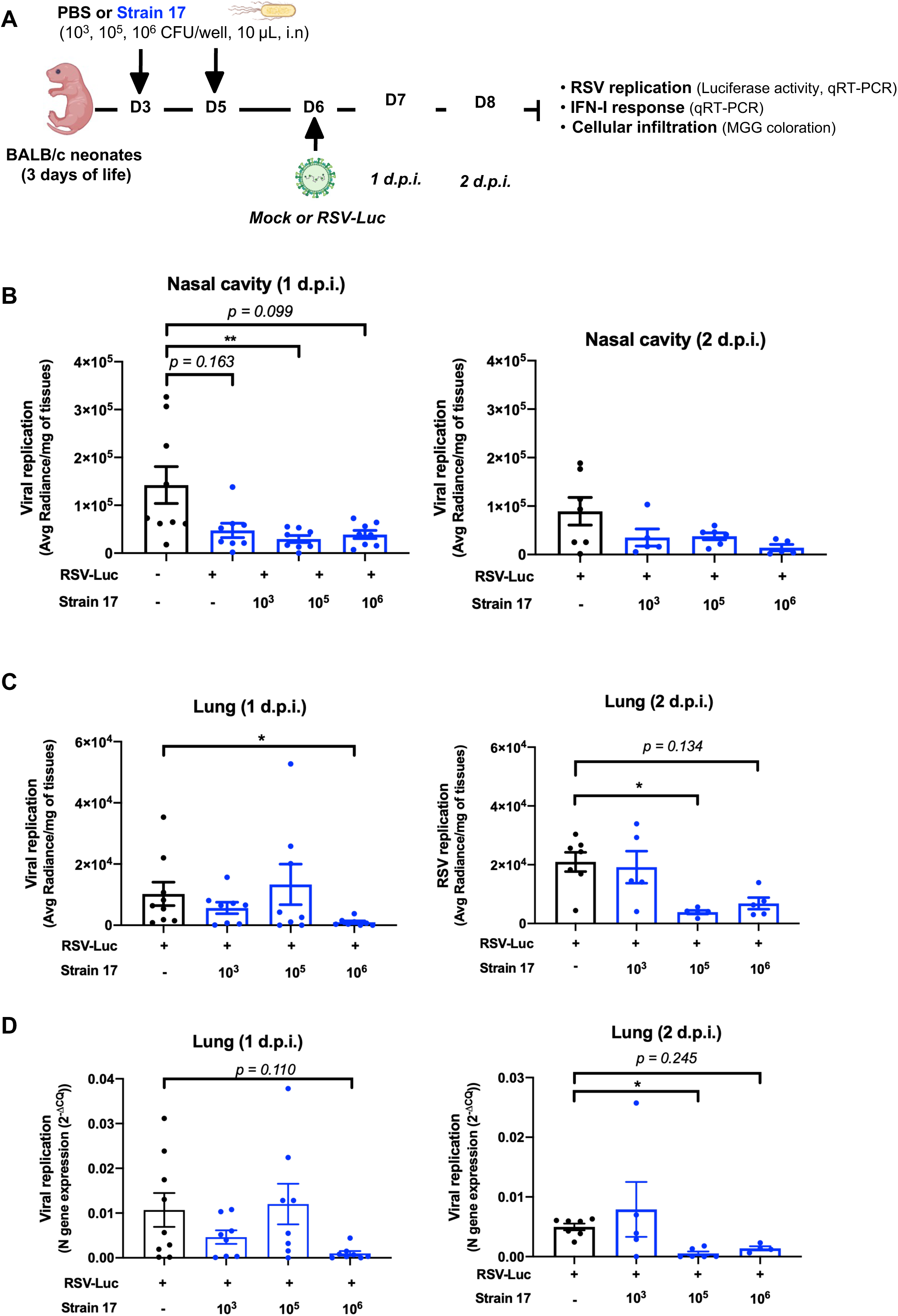
Early-life airway administration of strain 17 restricts RSV replication in neonatal mice. **A**. Experimental design. Neonatal mice were treated with different quantities of strain 17 (10^3^, 10^5^ or 10^6^ CFU) or PBS vehicle at 3 and 5-days old (i.n., 10 µL). At 6-days old, mice were infected with RSV-Luc (7.2 x 10^4^ PFU, 10 µL, i.n) and were euthanized at 1 d.p.i or 2 d.p.i. to assess viral replication by luciferase activity assay in the lysats of nasal cavities (**B**) and lungs (**C**). **D**. The viral replication was evaluated by quantification of the RSV *N gene* expression. The level of expression was calculated by the formula 2^-ΔCq^ with ΔCQ = Cq*_N gene_*-Cq*_reference genes_*. Results are expressed as mean ± SEM. Data are representative of two independent experiment (n = 5-8 per group). \**P* < 0.05, \*\**P* < 0.01 or adjusted *p*-value, ANOVA, Kruskal-Wallis test followed by Dunn’s post-hoc test.

**Figure. 5:**
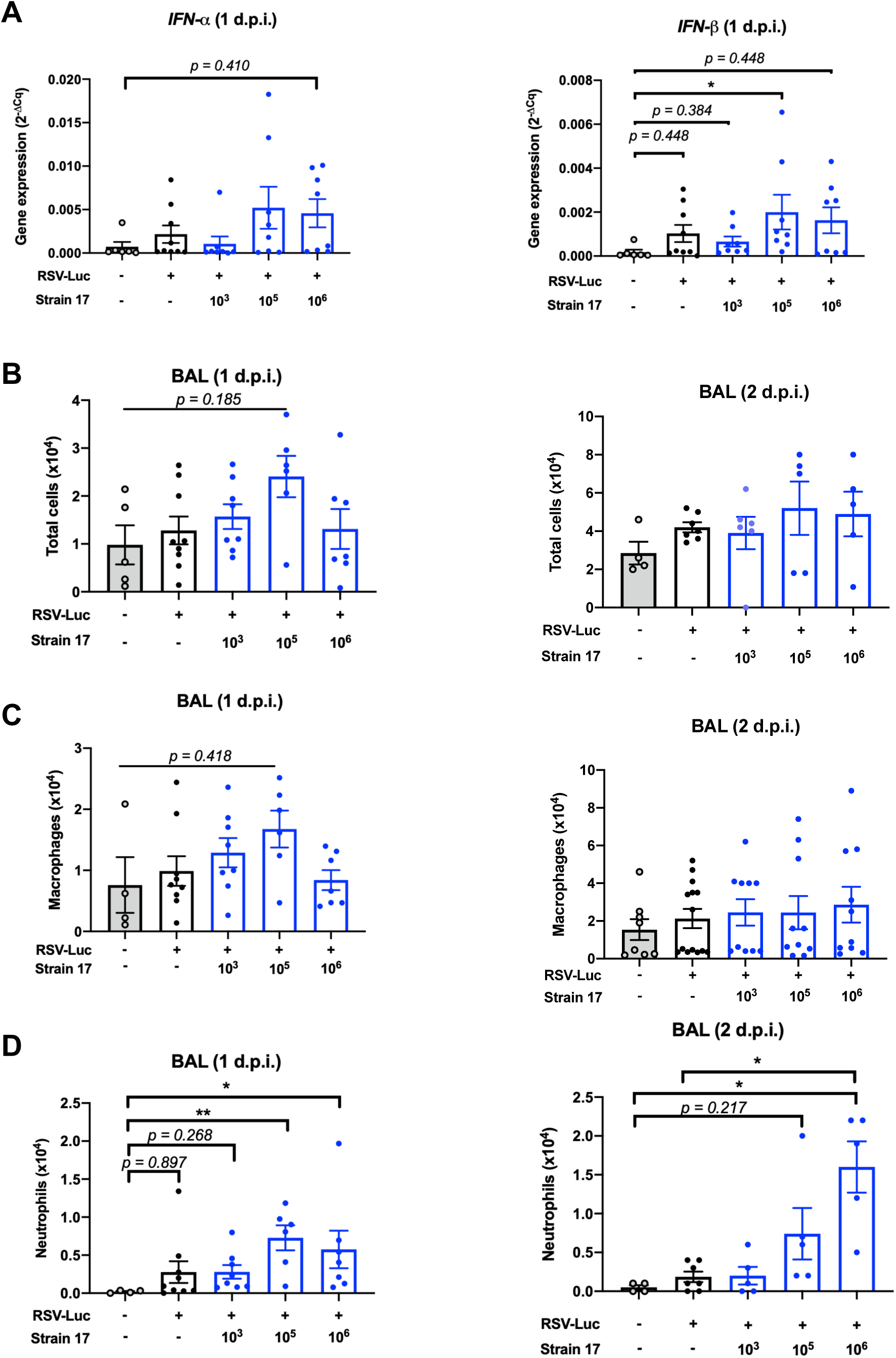
Early-life airway exposure to strain 17 enhances neonatal pulmonary immunity against RSV. Neonatal mice were treated with different quantities of strain 17 (10^3^, 10^5^ or 10^6^ CFU) or PBS vehicle at 3 and 5-days old (i.n., 10 µL). At 6 days old, mice were infected with RSV-Luc (7.2 x 10^4^ PFU, 10 µL, i.n) and were euthanized at 1 or 2 d.p.i. **A**. *IFN-I gene* expression was measured by qRT-PCR in lung lysates at 1 d.p.i.. Gene expression levels were calculated using the 2^-ΔCq^ method, with Cq values normalized to housekeeping genes. **B**. Cell counts were performed on BAL samples at 1 or 2 d.p.i.. Cell infiltration in BALs was assessed by May-Grünwald-Giemsa staining to identify and differentiate macrophages (**C**) and polynuclear neutrophils (**D**) at 1 or 2 d.p.i.. Results are expressed as mean ± SEM. Data are representative of two independent experiments (n = 5-8 per group). \**P* < 0.05, \*\**P* < 0.01 or adjusted *p-*value, ANOVA, Kruskal-Wallis test followed by Dunn’s post-hoc test.

### Neonatal strain 17 exposure reduces long-term pulmonary consequences of early-life RSV infection upon adult rechallenge

Exposure of neonatal PLCS, AMs, or lung tissues to strain 17 demonstrated a pronounced enhancement of type 1 immune responses. It has previously been demonstrated that enhancing type 1 response during neonatal primary RSV infection improves disease outcome upon secondary viral exposure. Thus, we employed a previously described model of adult RSV re-exposure to assess whether strain 17 exposure is associated with enhanced protective immunity and reduced immunopathological lung priming induced by early-life RSV infection [8]. Three-day-old mice were exposed intranasally with strain 17 or PBS vehicle at days 3, 5, 8, and 10 days of age. At 6-days-old, mice were infected intranasally with RSV, followed by a second exposure in adulthood (Fig 6A).

**Figure. 6:**
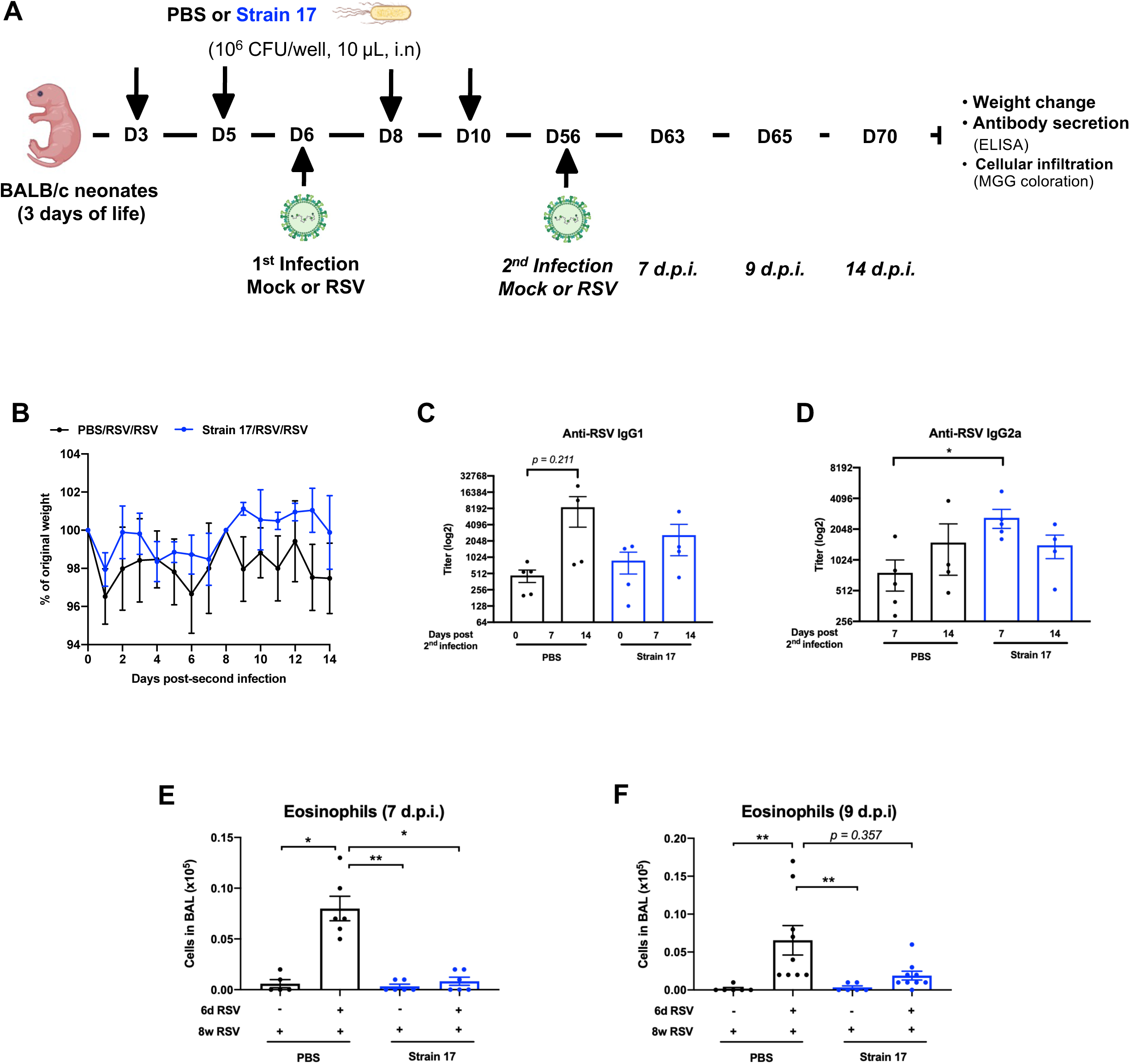
Early-life airway exposure to strain 17 attenuates the long-term pulmonary consequences of neonatal RSV infection during reinfection in adulthood. **A**. Experimental design. Three-day-old mice were treated intranasally with bacterium 17 (10⁶ CFU, i.n., 10 µL) or PBS vehicle at days 3, 5, 8, and 10 days of age. At 6 days old, mice were infected intranasally with RSV-A2 (2.3 × 10⁴ PFU in 10 µL), followed by a second infection with RSV-A2 (2.3 × 10⁵ PFU in 100 µL, i.n.) at 8 weeks of age. Mice were euthanized at 7, 9 or 14 days post-reinfection (d.p.i.) to assess cellular infiltration in the BALs and blood was collected. **B**. Weight evolution after the second infection. Results are expressed as mean ± SEM and presented as a percentage of the starting weight (n = 4 per group). **C-D**. Anti-RSV antibody assays. Serum was used to quantify anti-RSV IgG1 (**C**) or IgG2a (**D**) antibodies by ELISA. Results are expressed as the mean ± SEM (n = 4-5 per group). \**P* < 0.05, \*\**P* < 0.01, Mann-Whitney test. **E-F**. The infiltration of eosinophils in BALs was assessed by May-Grünwald-Giemsa staining at 7 or 9 d.p.i.. Results are expressed as mean ± SEM. Data are representative of two independent experiments (n = 5-9 per group). \**P* < 0.05, \*\**P* < 0.01 or adjusted *p-*value, ANOVA, Kruskal-Wallis test followed by Dunn’s post-hoc test.

Weight change was monitored (Fig 6B), and antibody responses (Fig 6C-D) and cell infiltration in BALs were assessed (S3B-E Fig). As expected, mice initially infected as pups and re-exposed to RSV in adulthood (PBS/RSV/RSV group) lost weight for over the 14 d.p.i., whereas mice administrated with strain 17 and infected with RSV as neonates (strain 17/RSV/RSV group) lost little weight and regained their initial weight by 8 d.p.i., consistently showing a better although not stastically significant, maintenance of body weight compared with controls (Fig 6B). We next investigated whether neonatal strain administration would impact the anti-RSV IgG1 response indicative of type 2 immunity and expected for adults upon RSV reexposure [8]. As expected, the re-exposure to RSV in adulhood induced a substantial production of anti-RSV-IgG1 at 14 d.p.i. in the PBS/RSV/RSV group (Fig 6C). In contrast, strain 17-exposed neonates exhibited early induction of anti-RSV-IgG2a antibodies, detectable as early as 7 d.p.i. (Fig 6D), with no significant increase in IgG1 levels at 14 d.p.i. (Fig 6C). Finally, we monitored cellular inflammation. At 7 days post-infection, total BAL cell numbers tended to be higher in both groups that were re-exposed to RSV as adults following a neonatal primary infection, whether the primary infection was with PBS/RSV/RSV (2.51 × 10⁵ ± 0.38 × 10⁵ cells) or with strain 17/RSV/RSV, compared to mice infected only as adults (PBS/Mock/RSV group, 1.26 × 10⁵ ± 0.33 × 10⁵ cells). However, this increase did not reach statistical significance relative to the PBS/Mock/RSV controls (S3B Fig). Likewise, no significant difference was observed between the PBS/RSV/RSV and strain 17/RSV/RSV groups (S3B Fig), indicating that neonatal priming with strain 17 did not significantly alter the magnitude of cellular inflammation in the BAL compared to a standard neonatal RSV priming. Macrophage and neutrophil numbers were similar across groups at 7 and 9 d.p.i. (S3C–D Fig), while the overall leukocyte increase was mainly due to lymphocyte recruitment (0.98 × 10⁵ ± 0.19 × 10⁵ cells in PBS/RSV/RSV *vs.* 0.81 × 10⁵ ± 0.24 × 10⁵ cells in strain 17/RSV/RSV; S3E Fig), which persisted until 9 d.p.i. despite declining over time. Significant eosinophil recruitment was observed in the BAL of PBS/RSV/RSV mice at 7 d.p.i. (0.08 × 10⁵ ± 0.012 × 10⁵ cells) and persisted at 9 d.p.i. (0.065 × 10⁵ ± 0.019 × 10⁵ cells), but was absent in PBS/Mock/RSV mice, reflecting type 2 immune priming from early-life RSV infection (Fig 6E–F). Notably, eosinophil infiltration was nearly abolished in mice exposed to strain 17 prior to the first RSV infection (0.006 × 10⁵ ± 0.004 × 10⁵ cells at 7 d.p.i.). Accordingly, the strain 17/RSV/RSV group had 9.6- and 3.4-fold fewer eosinophils in the BAL compared to PBS/RSV/RSV animals at 7 and 9 d.p.i., respectively. Neonatal exposure with strain 17 did not affect overall leukocyte infiltration upon adult RSV re-exposure but prevented immunopathologic eosinophilia. Altogether, these data indicate that the presence of strain 17 during primary RSV infection in early life was associated with a shift in the RSV-specific antibody response toward a type 1 profile (higher IgG2a and lower IgG1 levels), accompanied by minimal eosinophil recruitment and improved disease outcomes, as reflected by a trend toward better weight maintenance).

### Exposure to strain 17 is associated with reduced RSV spread in a translational human airway model

Finally, to assess whether strain 17 pre-exposure induces an antiviral state that limits subsequent RSV replication in a translational model, human primary airway epithelial cells cultured at the air-liquid interface were exposed to strain 17 and subsequently infected with mCherry-expressing RSV to visualize cellular RSV replication (Fig 7A). Without bacterial pre-exposure, RSV replication was detectable at 48h post-infection, as shown by mCherry fluorescence (Fig 7B). In contrast, strain 17 exposure markedly reduced mCherry activity compared to the control (Fig 7B-C). No cytotoxicity was detected (Fig 7D), and neither RSV exposure nor treatment with strain 17 induced the production of cytokines (IL-6, TNF-α or IP-10, S4 Fig) in these experimental conditions. Type III interferons (IFN-λ) are produced by airway epithelial cells infected with influenza virus earlier than IFN-I, helping to restrict viral spread without provoking inflammation [32]. As expected, RSV infection induced IFN-λ production in epithelial cells (Fig 7E). Strain 17 exposure alone did not induce IFN-λ and did not enhance the IFN-λ response to RSV infection; if anything, it tended to reduce secretion. Overall, the effect of strain 17 exposure is therefore unlikely to be explained by an increase in IFN-λ. We then measured another antiviral factor, the antimicrobial peptide human β-defensin 2, in epithelial cells (Fig 7F). This cationic molecule, secreted by lung epithelial cells, blocks RSV entry by destabilizing its envelope [33]. Strain 17 exposure strongly induced β-defensin 2 (2040 ± 571 pg/mL *vs* 65 ± 19 pg/mL in controls), whereas RSV alone triggered low levels (120 ± 32 pg/mL). Thus, pre-exposure to strain 17 markedly reduced RSV propagation in a human airway epithelium model, likely *via* enhanced β-defensin 2 secretion.

**Figure. 7:**
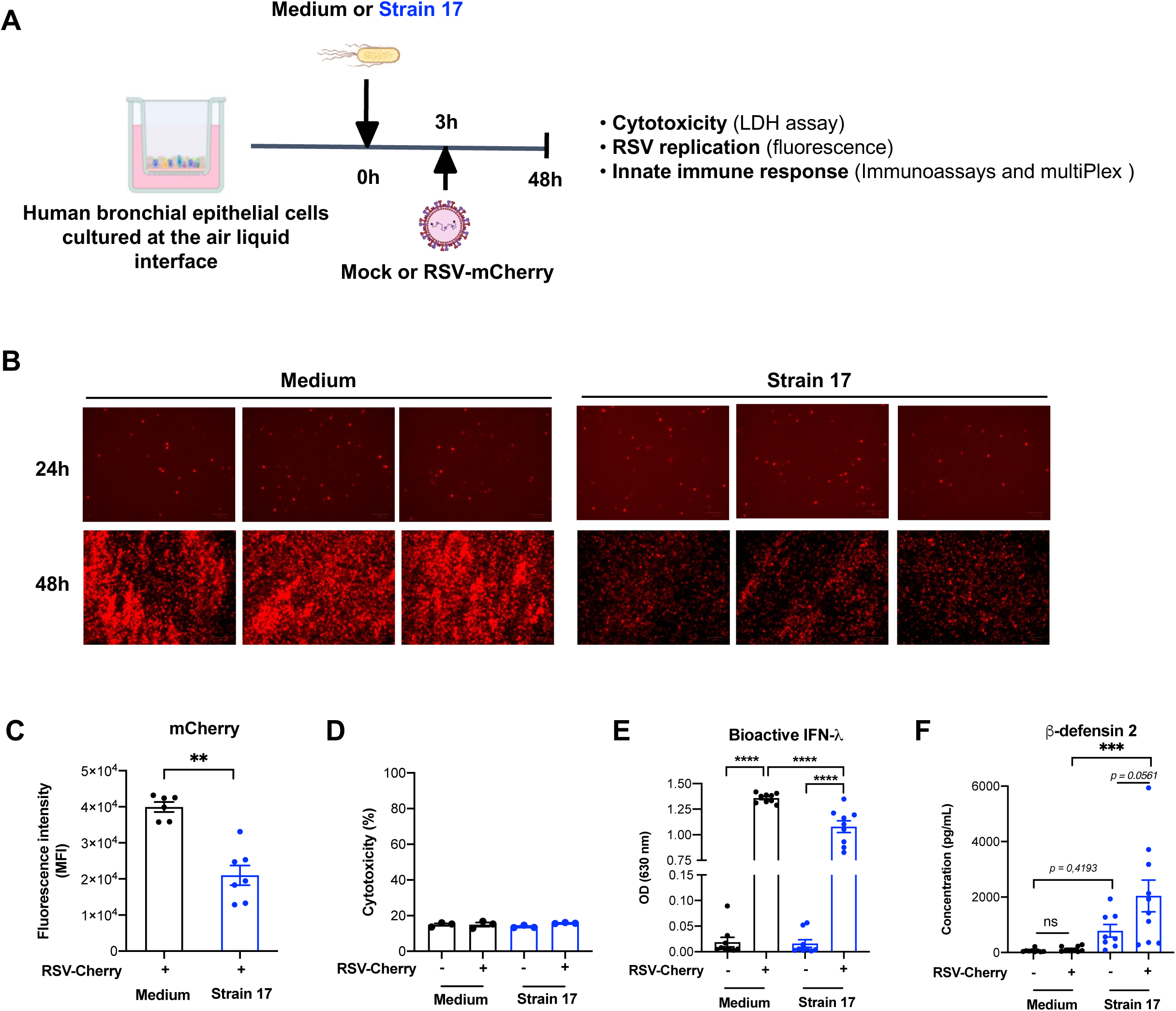
RSV replication in human primary airway epithelial cells is reduced by the pre-exposure to strain 17. **A**. Experimental design. Human primary bronchial epithelial cells cultured at the air liquid interface (MucilAir™) were first exposed to strain 17 (10^3^ CFU/well) for 3h, followed by RSV expressing mCherry (RSV-mCherry, MOI = 0.1) infection or an incubation with mock for a further 3h. Supernatants and cell lysates were collected 48h after incubation. RSV replication was evaluated by detecting mCherry using microscopy (**B**, three images representative of six inserts) or a fluorescence plate reader (**C**). Results are expressed as the mean fluorescence intensity (MFI) observed in lysats of RSV-mCherry-infected cells, after subtraction of the background MFI from mock-infected controls (± SEM), and represent a pool of two independent experiments (n = 3 inserts per group per experiment, \*\**P* < 0.01, Mann-Whitney test). **D**. Cytotoxicity observed in MucilAir™ was measured by LDH activity assay. Results are expressed as mean ± SEM. **E**. Bioactive IFN-λ was evaluated by measuring the response of HEK-Blue™ IFN-λ cells to the supernatants. SEAP activity was assessed using QUANTI-Blue™ Solution. The optical density (OD) at 630 nm is shown as mean ± SEM. **F**. β-defensin 2 secretion in supernatants was evaluated by ELISA. **E-F**. Data represent a pool of two independent experiments (n = 9 inserts per group, \*\*\**P* < 0.001, \*\*\*\**P* < 0.0001, 2way ANOVA with Tukey’s multiple comparisons test).

## Discussion

The lung microbiota and its influence on RSV infection severity have recently been identified as critical research areas in the fight against bronchiolitis in young children [24, 25] [22]. While modulation of the lung microbiota or metabolites appears to be a promising strategy for strengthening immune defenses in adults [34, 35], the impact during the critical period of lung mucosal development and susceptibility to RSV infection is yet unknown [29]. In this study, we demonstrated that the immune responses of neonatal mice to RSV infection, as well as their long-term consequences, could be favorably modulated by lung primo-colonizing bacterial strains given shortly after birth. In particular, using complementary cellular and *in vivo* approaches, we identified a bacterial strain 17 (*E. coli*), whose exposure promoted protective type 1 immune responses associated with reduced RSV replication in both mouse neonatal models and human epithelial cells.

We employed a PCLS-based screening platform from neonatal mice to assess cytokine responses to a collection of primo-colonizing bacterial strains isolated from mouse neonatal lungs. This strain collection encompasses major bacterial families, ranging from *Staphylococcus* to *Lactobacillus*. While most *Lactobacillus* strains were associated with low levels of immune activation in neonatal PCLS, and the majority of *Staphylococcus* strains with a pro-inflammatory cytokine profile, exposure to strain 17, identified as *Escherichia coli*, was associated with a distinct type 1-oriented cytokine response characterized by elevated levels of IFN-γ and IL-12p70 in the absence of detectable cytotoxicity. This immune profile is consistent with mechanisms involved in antiviral responses and may support enhanced antiviral immunity in the neonatal lung. However, exposure to strain 17 was also associated with increased production of IL-9, a cytokine linked to asthma and detected in airway aspirates from patients with severe bronchiolitis [36]. Its role in early-life pulmonary mucosa and RSV pathology remains poorly understood [37]. Interestingly, strain 17 exposure also induced the secretion of IL-22, which is known to promote epithelial barrier regeneration [38], and has a beneficial inhibitory effect on RSV infection has been demonstrated in a neonatal mouse model [39]. Although cytokine accumulation during 24-hour PCLS-based screening prevents the evaluation of secretion kinetics or autocrine effects, this approach nevertheless reveals the broad immunological responses associated with exposure to early-life lung primo-colonizing bacteria, including distinct patterns of immune stimulation and polarization. We acknowledge that bacterial biomass may differ between strains at the end of the 24-hour co-culture period, and that this could contribute to the observed differences in cytokine profiles. However, the primary objective of this study was not to isolate strain-specific effects independently of growth dynamics, but rather to demonstrate that early-colonizing strains induce distinct airway epithelial responses under standardized initial conditions. To this end, all strains were inoculated at the same MOI and cultures were grown to the early stationary phase prior to exposure, ensuring comparable physiological states at the time of inoculation. The differences in cytokine profiles observed across strains therefore reflect their overall immunostimulatory potential under biologically relevant conditions. Strain 17 differed markedly from the other lung primo-colonizing strains tested, being the only strain associated with reduced RSV replication in both PCLS and a human airway epithelium model, highlighting the relevance of further investigating its interactions with the host immune response. It is interesting to note that strains taken individually may display disease-specific immunobeneficial effects. Indeed, strain 24, previously shown to be protective in a juvenile asthma model [20], had neither a protective nor a detrimental effect on RSV replication in this study. These findings underscore the diverse immunomodulatory potential of bacteria within the pulmonary microbiota. In human epithelial cells, the reduced RSV replication observed following exposure to strain 17 appeared to be independent of IFN-λ and was associated with increased production of β-defensin 2, a host defense peptide known to destabilize viral envelopes and potentially contribute to limiting RSV infection and/or propagation [33]. Our study also demonstrated that strain 17 exposure can prime anti-RSV immunity in neonatal AMs by enhancing their production of IFN-I. Taken together, these findings suggest that exposure to strain 17 may contribute to the establishment of protective innate immune responses at the lung mucosa during early life.

Our *in vivo* studies confirmed the exposure of strain 17 to reduce RSV replication in the upper and lower respiratory tracts of neonatal mice. The degree of protection was dose-dependent and associated with an induction of IFN-I expression, as well as increased neutrophil recruitment. Moreover, early-life exposure to strain 17 mitigated long-term immunopathology upon re-exposure to RSV in adulthood. Mice treated with strain 17 exhibited a relatively stable body weight curve with no major alterations following viral reinfection, a shift toward type 1 immunity (increased IgG2a, decreased IgG1), and reduced eosinophilic infiltration.The role of the neutrophils recruited in strain 17-treated mice, with or without RSV infection, remains unclear, particularly regarding their protective *versus* pathological effects in neonates [40]. While neutrophils are classically associated with pro-inflammatory responses, recent evidence suggests that specific bacterial exposures can shape their phenotype toward a protective role [41]. Continuous inhalation of inactivated *A. lwoffii* F78 has been shown to induce PDL1⁺ immature neutrophils in the lung, reducing Th2-driven pathology following early-life RSV infection [41]. Consistent with this, our data show that early-life exposure of strain 17 induces neutrophil recruitment and confers lasting protection against RSV reinfection in adulthood, suggesting that bacterially-driven neutrophil responses may contribute to long-term consequences of type 2 immune imprinting established by neonatal RSV infection. Although colonization was not assessed, our study highlights the importance of dosing relative to the targeted respiratory region. For instance, the instillation of 10⁶ CFU into the lungs of a neonatal mouse represents a substantial bacterial load compared to the naturally sparse neonatal lung microbiota [20]. Higher doses were required to confer early protection in the lungs, whereas lower doses sufficed for the upper airways. This likely reflects the larger tissue surface of the lungs that must be exposed to the bacteria, rather than differences in microbiota density, which is already low in the pulmonary mucosa [42]. However, before proposing this bacterial strain as a therapeutic candidate for early immunomodulation to prevent acute and chronic RSV-related diseases, it is essential to elucidate the molecular pathways underlying its immunostimulatory effects in both epithelial and immune cells. Future studies should investigate whether the beneficial effects of strain 17 require live microorganisms, which colonize tissues and are likely more effective than inert components to produce metabolites. Testing conditioned supernatants could then clarify the contribution of bacterial metabolic activity, such as acetate production *in vitro* (data not shown), a metabolite known to enhance antiviral responses in AMs during RSV infection [43, 44]. Indeed, in an adult context, airway administration of *Lactobacillus* or *Corynebacterium* strains [45, 46], or a standardized bacterial lysate derived from human respiratory strains [47] is associated with improved resistance of mice to RSV challenge. Morever, higher acetate levels boosts the IFN-β response of AMs and epithelial ISG expression, reducing viral load and pulmonary inflammation in RSV-infected adult mice [43, 44]. It has been shown that *E. coli* probiotic strains, can have both deleterious and beneficial effects on digestive health [48]. So, before exploiting a such promising strain as a probiotic to enhance respiratory defenses, a global safety assessement is required.

Overall, this study provides a proof of concept that the immune responses of neonatal mice to RSV infection, as well as their long-term consequences, can be favorably modulated by bacterial strains that colonize the lung early in life. These findings further support the notion that the neonatal lung microbiota plays a key role in shaping pulmonary immune responses, and that its disruption, for example through antibiotic use, can have profound and lasting effects on host immunity and respiratory vulnerability [49]. In line with this, a recent study in pediatric RSV cohorts indicates that persistent respiratory symptoms during the convalescence phase are associated with disruptions of the respiratory microbiota [26]. It would be interesting to further explore the potential benefits of primo-colonizing strains during recovery, opening new perspectives for preventing adverse lung trajectories through early-life microbiota modulation.

## Materials and methods

### Bacterial strains’ identification and growth

Bacterial strains (S1 Table 1) previously isolated from 3-day-old mouse lungs were stored at −80°C [20]. Strains were revived in 10 mL yeast-heminBHI or MRS medium and subcultured (100 µL/10 mL). Growth was monitored hourly for 8h by spectrophotometry to determine exponential and stationary phases (data not shown). Bacterial counts were obtained using a BD Accuri™ C6 flow cytometer, and strain identity was confirmed by 16S PCR (J4/J7 primers) and MALDI-TOF mass spectrometry (CIRM, INRAE, Nouzilly, France).

**Supplemental table S1:**
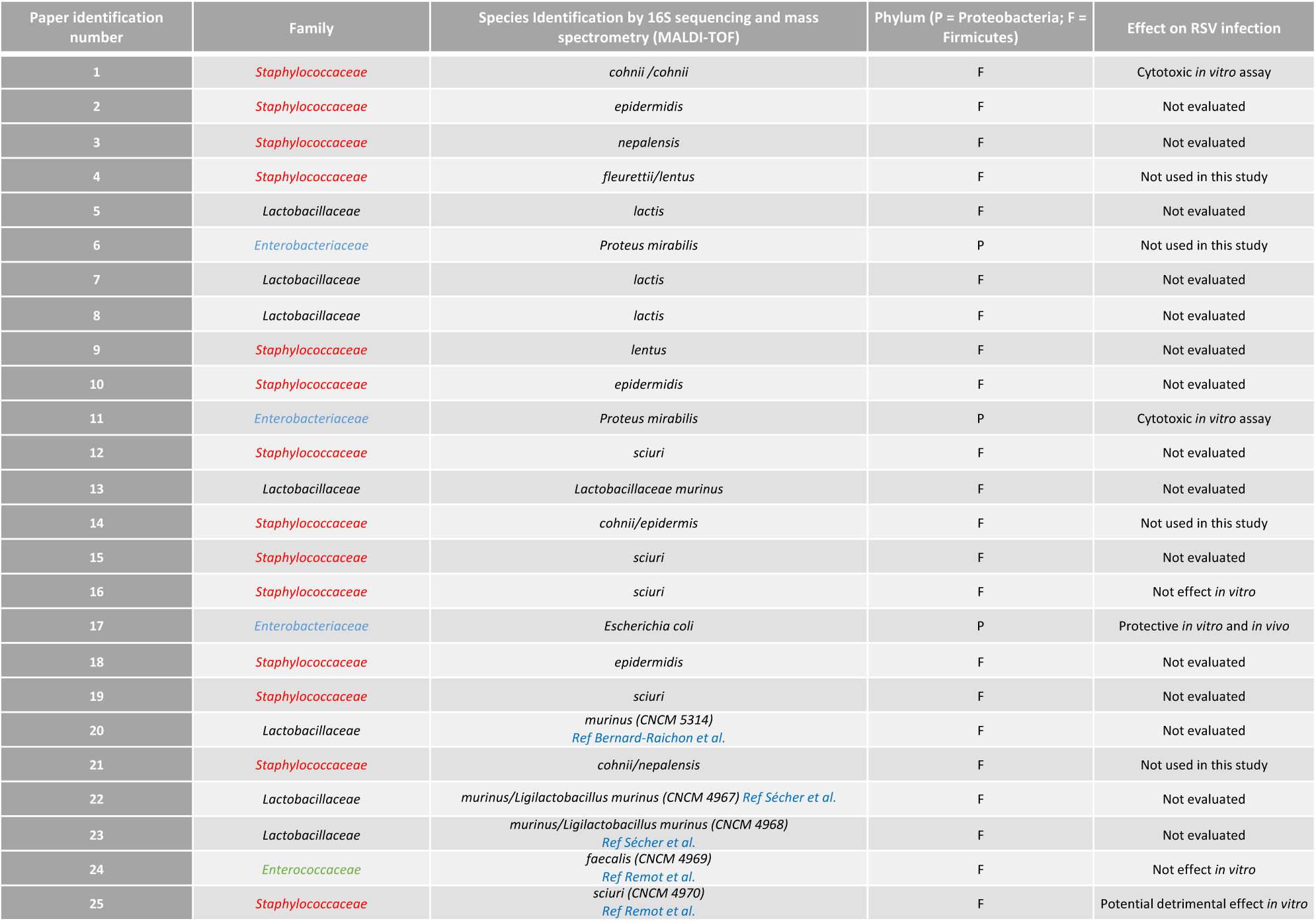
Descriptive table of lung primo-colonizing commensal bacteria isolated from neonatal mice.

### Virus

Human RSV strain A2 and recombinant RSV-Long expressing mCherry (RSV-mCherry) or luciferase (RSV-Luc) were produced, titrated on HEp-2 cells, and stored at -80°C [50].

### Ethics Statement

All experiments were approved by the ethics committee COMETHEA (Ethical Committee for Animal Experimentation, authorization number: APAFIS#14654-2018032116445169). Pregnant BALB/cJrj mice were purchased from Janvier (Le Genest, France).

### *Ex vivo* lung explants, stimulation and infection

Lungs from 6-day-old neonatal mice were filled *via* tracheotomy with RPMI containing 1.5% low-melting agarose (37°C). After solidification, lungs were placed in a Krumdieck tissue slicer (MD6000) chamber containing PBS with 1× antibiotic-antimycotic and cut into 150 µm slices to obtain precision-cut lung slices (PCLS) [20, 22]. After plating, the PCLS were washed with antibiotic-free medium and incubated overnight at 37 °C. PCLS were stimulated for 16h at 37 °C, 5% CO₂ with strains (50 CFU/well in antibiotic-free RPMI medium with 5% FBS and L-glutamine and without antibiotic). Inoculations were verified by CFU counts on agar plates. Strains were standardized at the onset of the stationary phase to ensure a comparable bacterial physiological state at the time of stimulation. Three hours later, PCLS were infected with RSV-Luc (7.19 × 10⁴ PFU/well) or Mock control. Supernatants were collected. PCLS were lysated in lysis buffer (5 mM EDTA, 150 mM NaCl, 50 mM Tris-HCl, 1% Triton X100, pH 7,4) or Passive Lysis Buffer (30 mM Tris, 10 mM MgCl_2_, 1% Triton 100X, 2% Glycerol) using a Precellys 24-bead grinder homogenizer (Bertin Technologies) for 2 × 15s at 6,000 x g. Protein levels were quantified with DC™ or RC-DC™ Protein Assays (Bio-Rad).

### Experiments with differentiated human primary bronchial epithelial cells

MucilAir^®^ from healthy donors were purchased from Epithelix. Cells (≈5 x 10^5^ cells/transwell) were stimulated with bacterium 17 (10^3^ CFU/transwell) in 20 µL in apical side. After 3h, cells were infected with RSV-mCherry (MOI = 0.1) or Mock control during 3h. The inoculum was removed and medium replaced by strain 17 containing fresh medium. RSV-mCherry replication was visualized using the ZOE Fluorescent Cell Imager (Bio-Rad) and quantified with a fluorescence plate reader (Tecan Infinite 200 Pro). Supernatants and cell lysates were then collected.

### *Ex vivo* AM culture and infection

Bronchoalveolar lavages (BALs) were performed on 6-days-old mice to isolate AM, as previously described [9, 51, 52]. Cells (1 x 10^5^) were plated, 24h after seeding, incubated with bacterial strains (50 CFU/well) for 3h before exposure to RSV-Long (MOI = 5) or a mock control. Supernatants and cells were collected 24h post-infection.

### Cell cytotoxicity

Cytotoxicity was measured by lactate deshydrogenase (LDH) activity, using the Non-Radioactive Cytotoxicity Assay kit (Promega).

### Cytokine, IFN, and RSV-specific Ab assays

Cytokine production was measured using the 15-plex or IL-6/TNF-α/IFN-α/IFN-β custom 4-Plex mouse ProcartaPlex™ immunoassay (ThermoFisher Scientific). Human IL-6, TNF-α or IP-10 were measured by Luminex and human β-defensin-2 by ELISA (NB-50-1634-1000, NeoBiotech). Data were read on MagPix instrument (Luminex), and analyzed using Bio-Plex Manager software (Bio-Rad). HEK-Blue™ IFN-λ (InvivoGen) cells were used for the detection of bioactive human IFN-λ. Individual mouse sera were assayed for specific Ab (IgG1, IgG2a) against RSV lysates by ELISA [13].

### *In vivo* bacterial strains inoculation and RSV infection

Neonate mice (age 3 and 5 days) received 10 µL of strain 17 intranasally (i.n.) route. At 6 days old, mice were infected with RSV-Luc (7.187 x 10^6^ PFU/mL, 10 µL, i.n) or a Mock control. In other experiments, neonate mice were first infected by RSV-A2 (2.3 × 10^6^ PFU/mL, 10 µL, i.n.), and were kept until 8 weeks of age when they were all infected with RSV-A2 (100 µL, i.n.) [13]. BAL and lung tissues were collected.

### Bioluminescence measurements

Photon emission was measured in lysates of the nasal cavity or lung tissues and PCLS using the IVIS system (Xenogen Biosciences) and Living Image software (version 4.0, Caliper Life Sciences), following a previously described method [9], and was normalized to protein content.

### Cell infiltration

Cells were collected from BALs and centrifugated at 400 x g for 10min at 4°C. May-Grunwald-Giemsa coloration was performed to assess the differential cell counts using light microscopy. Two hundred cells were counted to determine relative and absolute cell numbers.

### RT-qPCR

Total RNA was extracted from lung lysates using a NucleoSpin® RNA midi column (Macherey Nagel) and reverse transcribed using random primers and SuperScript II (Invitrogen), according to the manufacturer’s instructions. The primers (Sigma–Aldrich) used are listed (S2 Table). RT-qPCR was performed in duplicate for each gene using the CFX Connect™ Real-Time PCR Detection System (Bio-rad) and SYBRGreen PCR Master Mix (Eurogenetec). Data were analyzed using CFX Manager™ Software (Bio-rad) to determine the quantification cycle (Cq) values. Results were determined using the formula 2^-ΔCq^, with ΔCQ = Cq_gene_- Cq_(mean (*bActine*,*Gapdh*, *Hprt*))_.

**Supplemental table S2:**
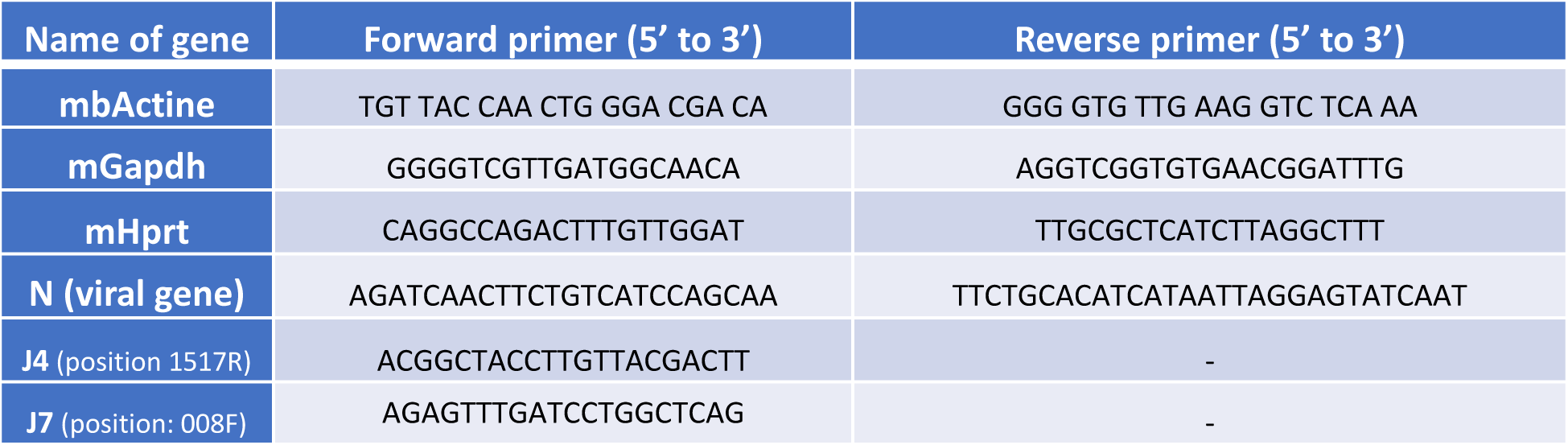
List of primers used for RT-qPCR.

### Data and statistical analysis

Data are expressed as mean ± SEM. Statistical significance was set at \**P* < 0.05, \*\**P* < 0.01, \*\*\**P* < 0.001, and \*\*\*\**P* < 0.0001 using GraphPad Prism 8 (GraphPad Software Inc.). Data normality was assessed using the D’Agostino–Pearson and Shapiro–Wilk tests. Depending on the distribution, differences between groups were analyzed using a parametric *t*-test or a nonparametric Mann–Whitney test (two groups), or by one-way ANOVA/Kruskal–Wallis or two-way ANOVA with Tukey’s multiple comparisons test.

## Acknowledgements

We greatly acknowledge M. Prost and A. Mialet for their assistances with experiments. We are grateful to M-A. Rameix-Welti (INSERM, U1173, UVSQ) and J-F Eléouët (INRAE, UVSQ, VIM) for providing recombinant RSV-Cherry. We are grateful to the IERP unit, INRAE (Infectiology of fishes and rodent facility, doi: 10.15454/1.5572427140471238E12). We thank the Emerg’in platform for access to IVIS-Spectrum that was financed by the Region Ile De France (DIM One Health). We thank E Helloin and C Darrigo (CIRM, INRAE, Nouzilly) for bacterial strains identification by mass spectrometry.

## Supporting information

**Supplemental Figure. 1: Cytokines production of precision-cut lung slices (PCLS) from neonatal mice in response to exposure to lung primo-colonizing bacteria.** The lungs from 6-day-old mice were isolated and cut with a Krumdieck tissue slicer to obtain PCLS. They were then exposed to bacteria (50 CFU/well) or cultured in control medium. Supernatants were collected 16h after stimulation with bacterial strains or PBS, and cytokine production was assessed using the 15-plex Mouse ProcartaPlex™ immunoassay. **A**. Cytotoxicity observed in PCLS after bacterial exposure was measured by LDH activity assay. **B**. Pro-inflammatory cytokine secretion (GM-CSF, IL-6, IL-1β, TNF-α). **C**. Cytokine associated with type 1 immune response (IL-18). **D**. Cytokines associated with type 2 immune response (IL-4, IL-5, IL-10, IL-13). **E**. Cytokines associated with type 17/22/9 immune response (IL-17A, IL-22, IL-9). **F**. Secretion of IL-2 associated with cell proliferation response. Results are expressed as mean ± SEM and represent a pool of five independent experiments (n = 2-5 biological replicate samples/condition). \**P* < 0.05, \*\**P* < 0.01, \*\*\**P* < 0.001, and \*\*\*\**P* < 0.0001 between the control medium and the bacteria-exposed condition (ANOVA, Kruskal-Wallis test followed by Dunn’s post-hoc test).

**Supplemental Figure. 2: Bronchoalveolar cellular response to intranasal exposure to strain 17 in neonatal mice. A.** Experimental design. Three-day-old mice were treated intranasally with strain 17 (10⁶ CFU, i.n., 10 µL) or PBS vehicle on days 3 and 5 of age. At day 7 and 8 of age, mice were euthanized and BAL were collected. **B**. Total cell counts in BAL samples. Cellular composition in BAL was assessed by May-Grünwald-Giemsa staining to identify and differentiate macrophages (**C**) and neutrophils (**D**). Results are expressed as mean ± SEM (n = 3 per group). \**P* < 0.05 between the PBS- and the bacteria-exposed condition (ANOVA, Kruskal-Wallis test followed by Dunn’s post-hoc test).

**Supplemental Figure. 3: Early-life airway delivery of strain 17 prevents weight loss during RSV reinfection in adulthood and shapes the adaptive antibody response. A.** Experimental design. Three-day-old mice were treated intranasally with strain 17 (10⁶ CFU, i.n., 10 µL) or PBS vehicle at days 3, 5, 8, and 10 of age. On postnatal day 6, mice were infected intranasally with RSV-A2 (2.3 × 10⁴ PFU in 10 µL), followed by a second infection with RSV-A2 (2.3 × 10⁵ PFU in 100 µL, i.n.) at 8 weeks of age. **B-E.** Mice were euthanized at 7 or 9 d.p.i. to assess cell infiltration in the bronchoalveolar lavage (BAL). **B**. Total cell counts in BAL. Cell infiltration in BAL was assessed by May-Grünwald-Giemsa staining to identify and differentiate macrophages (**C**), neutrophils (**D**), and lymphocytes (**E**) at 7 or 9 d.p.i.. Results are expressed as mean ± SEM. Data are representative of two independent experiments (n = 5-9 per group, addjusted *p-*value, ANOVA, Kruskal-Wallis test followed by Dunn’s post-hoc test).

**Supplemental Figure. 4: Cytokine response to RSV infection in primary human airway cells with or without strain 17 pre-exposure.** Human primary bronchial epithelial cells cultured at the air-liquid interface (MucilAir™) were first exposed to strain 17 (10^3^ CFU/well) for 3h, followed by RSV expressing mCherry (RSV-mCherry, MOI = 0.1) infection or incubation with mock. Basal medium and cell lysates were collected 48h after incubation. IL-6, TNF-α or IP-10 concentrations in basal medium were assessed using Luminex immunoassay. Means ± SEM are represented.

## Author Contibutions

Q.M. and C.C.: Methodology, Validation, Investigation, Data curation, Visualization, Writing – review & editing. C.F.: Validation, Data curation, Resources. E.P. and E.B.: Validation, Data curation. V.S-C. and D.L.: Methodology, Writing – review & editing. C.D.: Methodology, Investigation. A.R.: Methology, Conceptualization. S.C., S.H. and E.S.: Resources. I.S-C., M.T. and S.R.: Resources, Conceptualization, Writing – review & editing. D.D.: Conceptualization, Methodology, Investigation, Validation, Formal analysis, Writing – Original Draft, Writing – review & editing, Supervision, Visualization, Project administration, Funding acquisition.

## Data Availability Statement

Data are available from the corresponding author upon reasonable request.

