## Supplementary material for "Early-life exposure to commensal lung bacteria primes innate antiviral immunity and prevents RSV immunopathology in neonatal mice": Supp Figure

Supplemental Figures

### Supplemental Figure S1

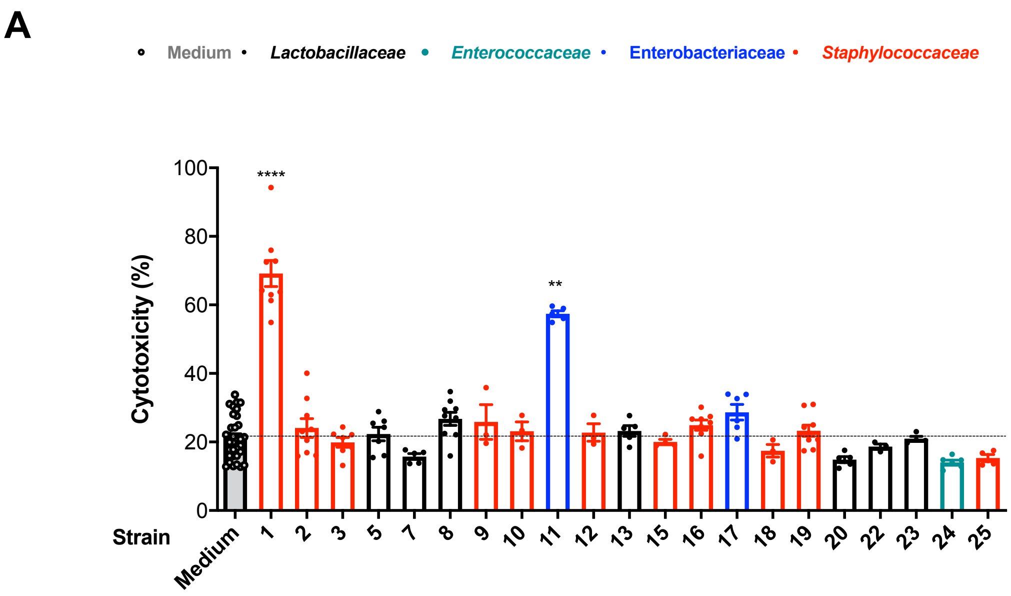

### Supplemental Figure S1

**B**

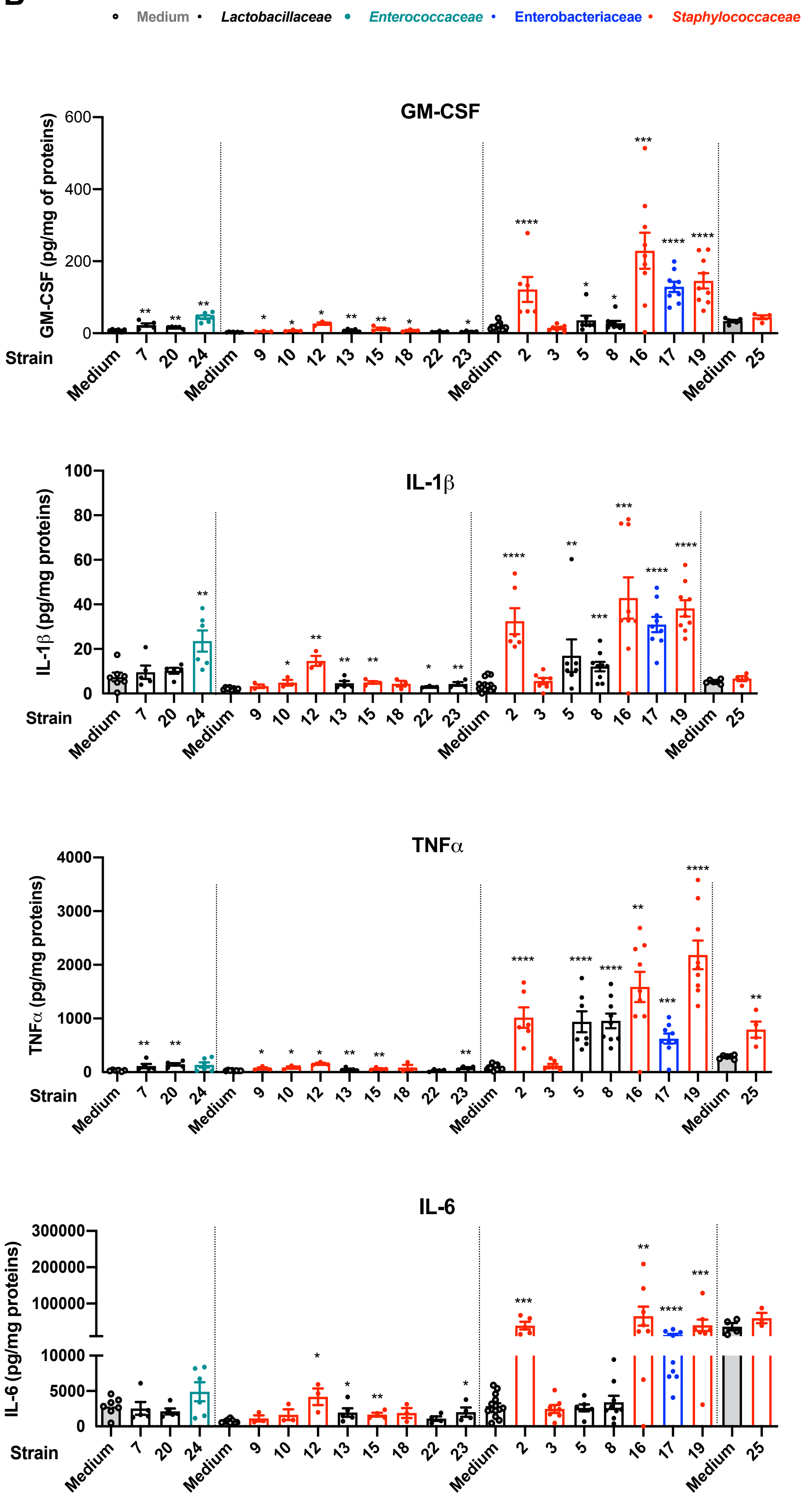

### Supplemental Figure S1

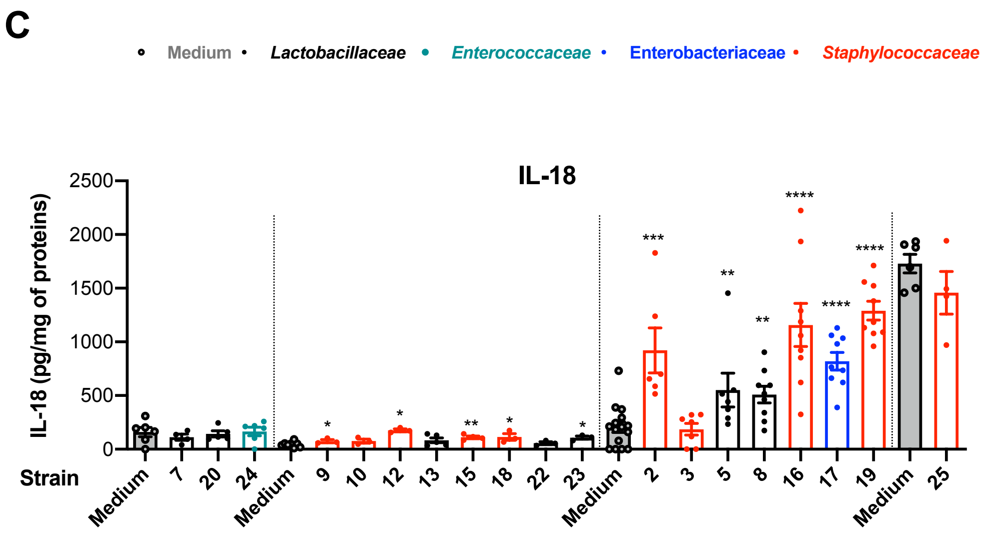

### Supplemental Figure S1

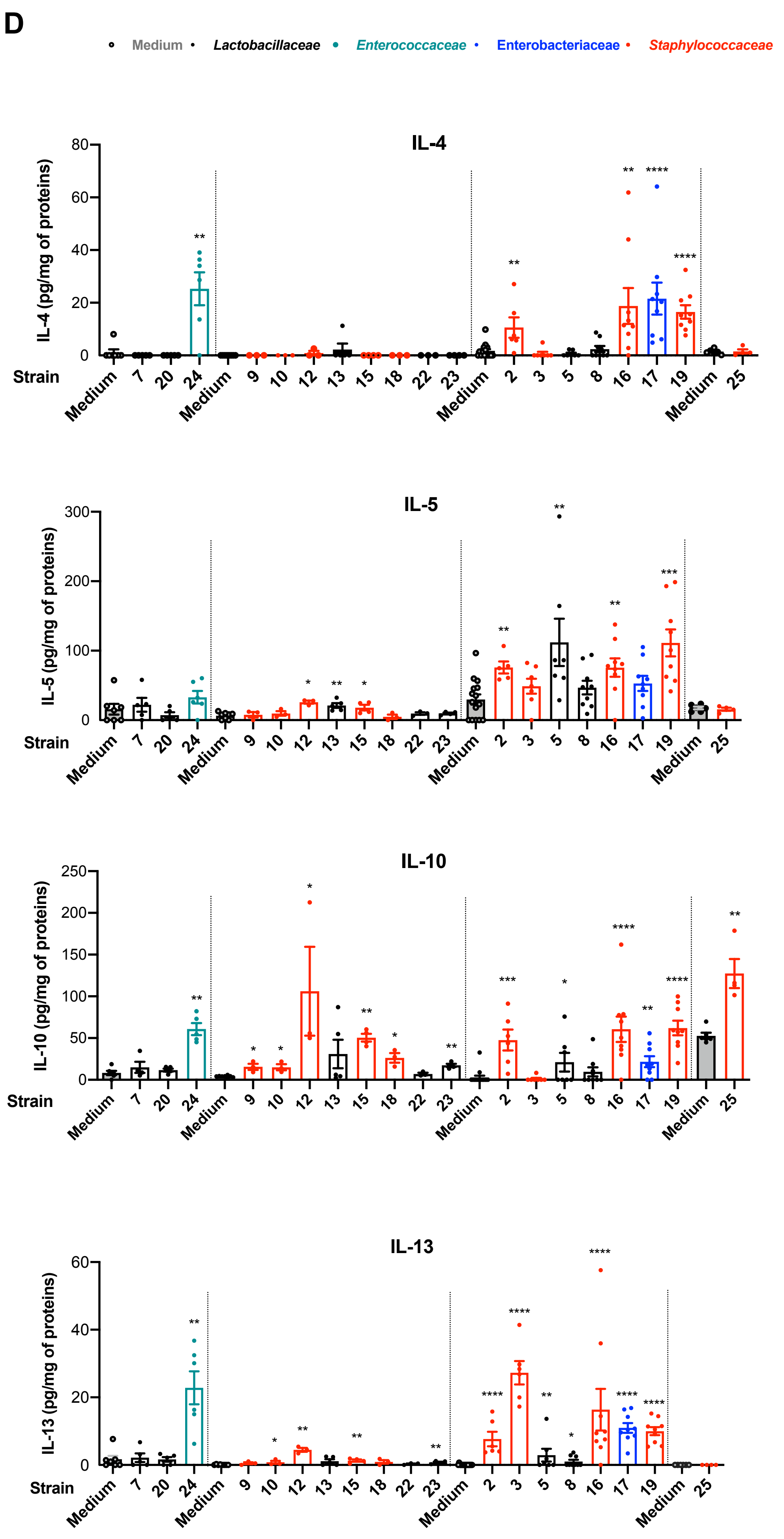

### Supplemental Figure S1

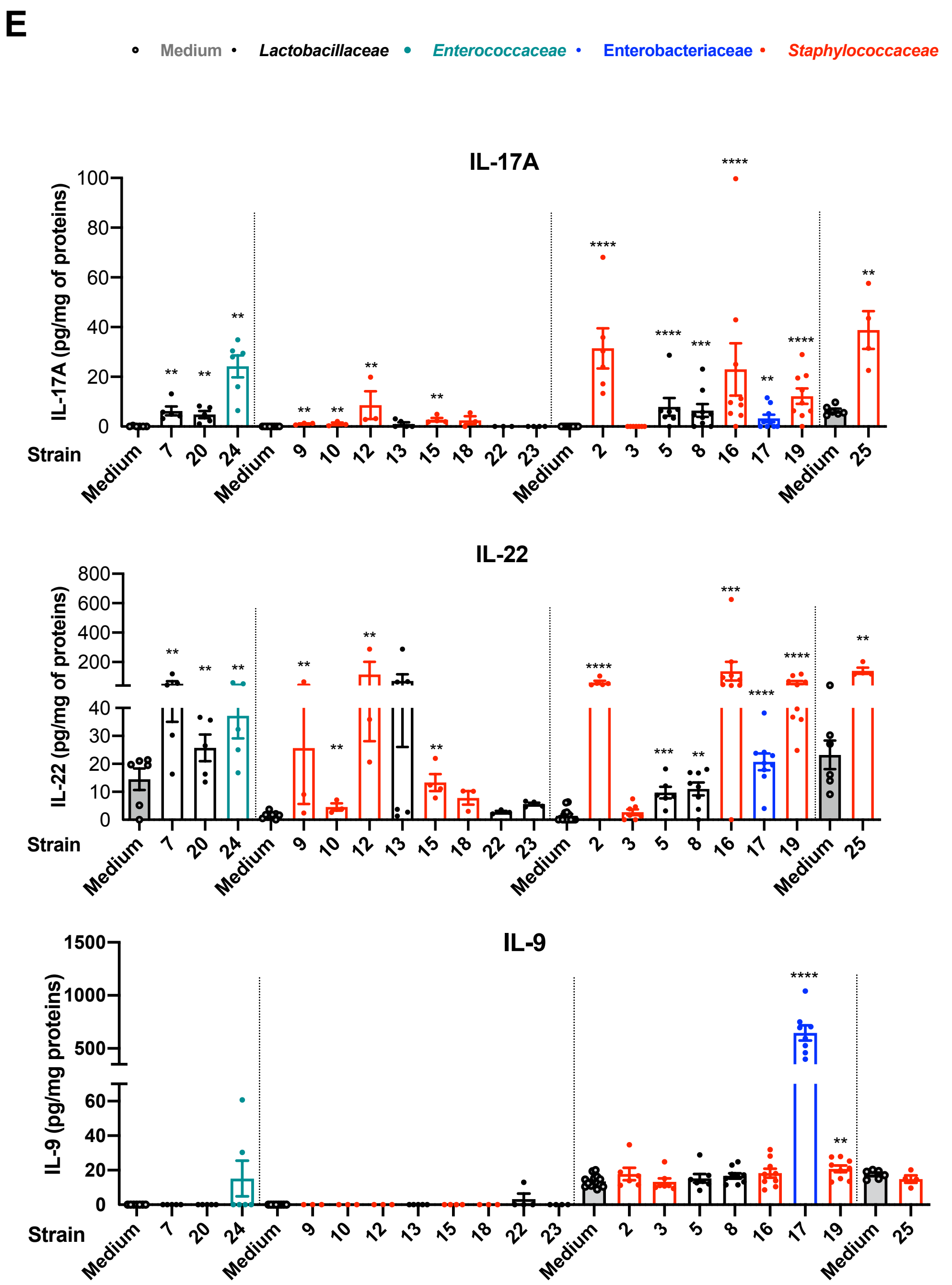


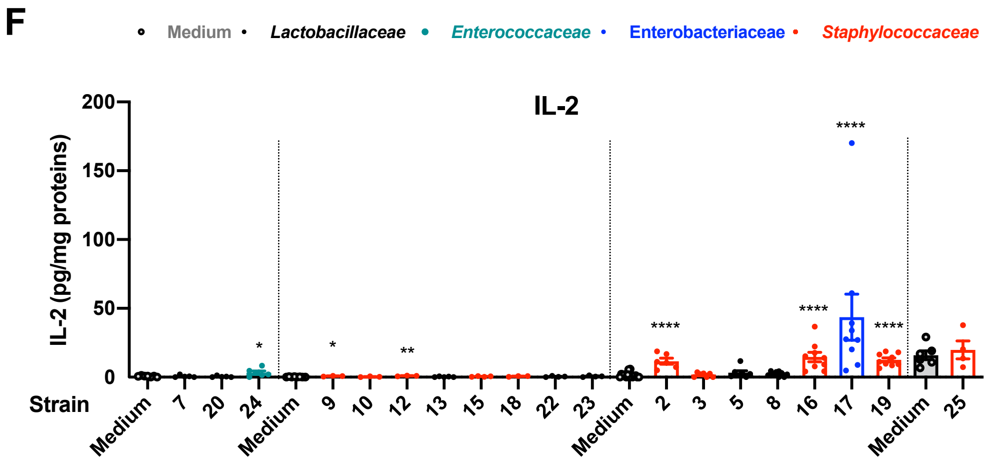

### Supplemental Figure S2

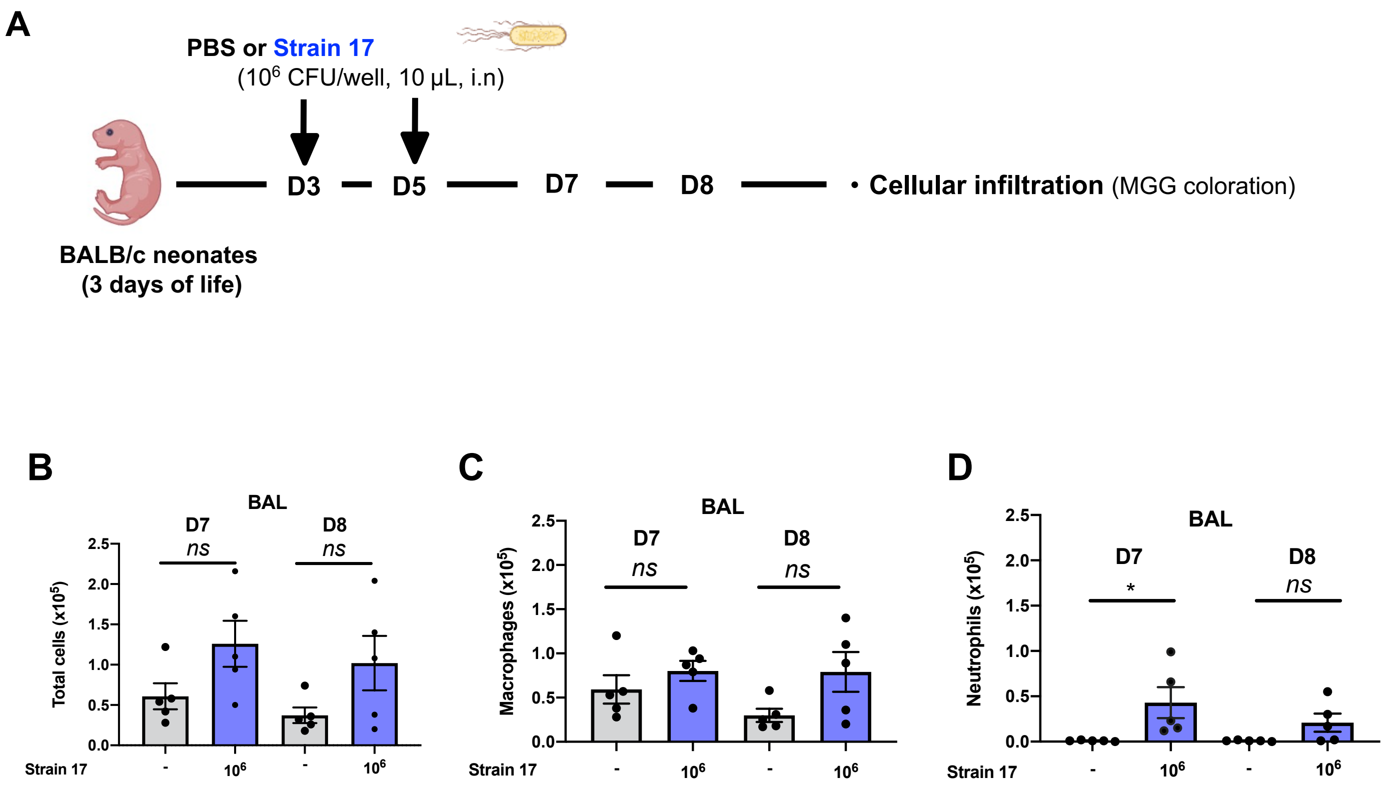

#### Supplemental Figure S3

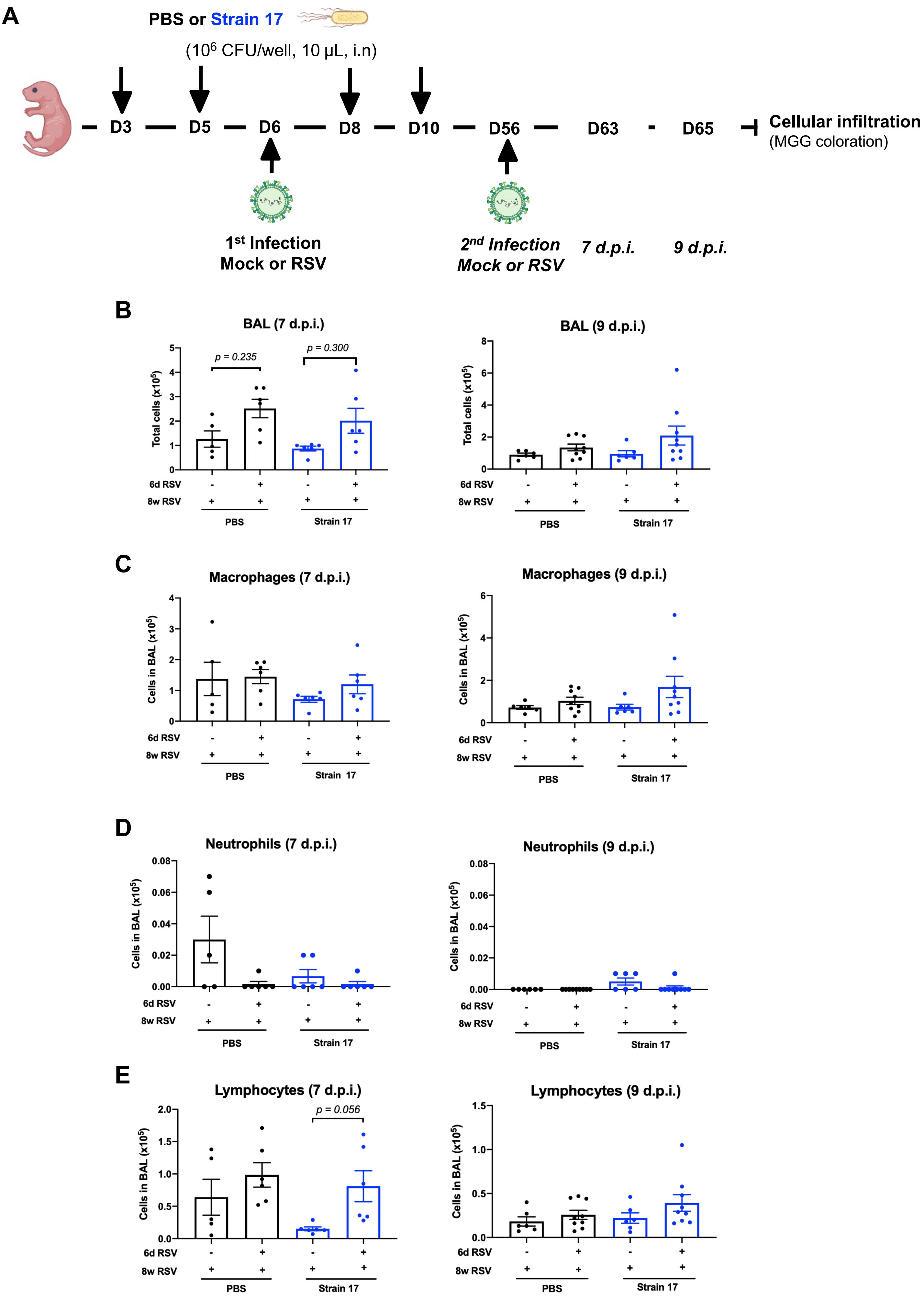

### Supplemental Figure S4

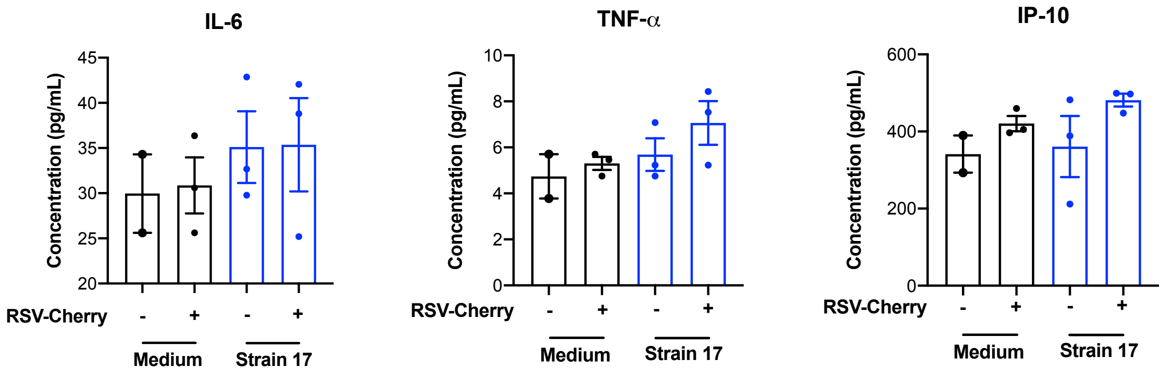
