## Supplementary material for "Early-life exposure to commensal lung bacteria primes innate antiviral immunity and prevents RSV immunopathology in neonatal mice": table

**Supplemental table S1**: Descriptive table of lung primo-colonizing commensal bacteria isolated from neonatal mice.

| Paper identification number | Family | Species Identification by 16S sequencing and mass spectrometry (MALDI-TOF) | Phylum (P = Proteobacteria; F = Firmicutes) | Effect on RSV infection |
|---|---|---|---|---|
| 1 | *Staphylococcaceae* | *cohnii /cohnii* | F | Cytotoxic *in vitro* assay |
| 2 | *Staphylococcaceae* | *epidermidis* | F | Not evaluated |
| 3 | *Staphylococcaceae* | *nepalensis* | F | Not evaluated |
| 4 | *Staphylococcaceae* | *fleurettii/lentus* | F | Not used in this study |
| 5 | *Lactobacillaceae* | *lactis* | F | Not evaluated |
| 6 | *Enterobacteriaceae* | *Proteus mirabilis* | P | Not used in this study |
| 7 | *Lactobacillaceae* | *lactis* | F | Not evaluated |
| 8 | *Lactobacillaceae* | *lactis* | F | Not evaluated |
| 9 | *Staphylococcaceae* | *lentus* | F | Not evaluated |
| 10 | *Staphylococcaceae* | *epidermidis* | F | Not evaluated |
| 11 | *Enterobacteriaceae* | *Proteus mirabilis* | P | Cytotoxic *in vitro* assay |
| 12 | *Staphylococcaceae* | *sciuri* | F | Not evaluated |
| 13 | *Lactobacillaceae* | *Lactobacillaceae murinus* | F | Not evaluated |
| 14 | *Staphylococcaceae* | *cohnii/epidermis* | F | Not used in this study |
| 15 | *Staphylococcaceae* | *sciuri* | F | Not evaluated |
| 16 | *Staphylococcaceae* | *sciuri* | F | Not effect *in vitro* |
| 17 | *Enterobacteriaceae* | *Escherichia coli* | P | Protective *in vitro* and *in vivo* |
| 18 | *Staphylococcaceae* | *epidermidis* | F | Not evaluated |
| 19 | *Staphylococcaceae* | *sciuri* | F | Not evaluated |
| 20 | *Lactobacillaceae* | *murinus (CNCM 5314)*<br>*Ref Bernard-Raichon et al.* | F | Not evaluated |
| 21 | *Staphylococcaceae* | *cohnii/nepalensis* | F | Not used in this study |
| 22 | *Lactobacillaceae* | *murinus/Ligilactobacillus murinus (CNCM 4967) Ref Sécher et al.* | F | Not evaluated |
| 23 | *Lactobacillaceae* | *murinus/Ligilactobacillus murinus (CNCM 4968)*<br>*Ref Sécher et al.* | F | Not evaluated |
| 24 | *Enterococcaceae* | *faecalis (CNCM 4969)*<br>*Ref Remot et al.* | F | Not effect *in vitro* |
| 25 | *Staphylococcaceae* | *sciuri (CNCM 4970)*<br>*Ref Remot et al.* | F | Potential detrimental effect *in vitro* |

**<u>Supplemental table S2</u>: List of primers used for RT-qPCR.**

| Name of gene | Forward primer (5' to 3') | Reverse primer (5' to 3') |
|---|---|---|
| **mbActine** | TGT TAC CAA CTG GGA CGA CA | GGG GTG TTG AAG GTC TCA AA |
| **mGapdh** | GGGGTCGTTGATGGCAACA | AGGTCGGTGTGAACGGATTTG |
| **mHprt** | CAGGCCAGACTTTGTTGGAT | TTGCGCTCATCTTAGGCTTT |
| **N (viral gene)** | AGATCAACTTCTGTCATCCAGCAA | TTCTGCACATCATAATTAGGAGTATCAAT |
| **J4** (position 1517R) | ACGGCTACCTTGTTACGACTT | - |
| **J7** (position: 008F) | AGAGTTTGATCCTGGCTCAG | - |
